# Replication-independent eviction of H2B.8 reveals chromatin reprogramming during seed imbibition

**DOI:** 10.1101/2025.08.21.670522

**Authors:** Lauriane Simon, Stefania Paltrinieri, Manon Verdier, Qingyi Wang, Sylviane Cotterell, David Latrasse, Aude Maugarny-Calès, Gilles Sireta, Sophie Desset, Simon Amiard, Christophe Bailly, Kentaro Tamura, Moussa Benhamed, Christophe Tatout, Samuel Le Goff, Aline V. Probst

## Abstract

The transition from seed to seedling involves major changes in nuclear organization and gene expression, yet the extent to which this developmental transition requires chromatin reprogramming remains largely unexplored. Here, we report that Arabidopsis dry seed embryos accumulate the histone variant H2B.8, which contributes to higher-order chromatin organization by forming spatial clusters that structure the 3D nuclear space. H2B.8 forms heterotypic nucleosomes at euchromatic transposons and lowly expressed genes and, during imbibition, modulates the transcriptional activation of a subset of these genes. Imbibition triggers a decrease in *H2B.8* transcripts and the eviction of H2B.8 proteins in a process that operates independently of DNA replication but requires protein translation and *de novo* transcription. Histone eviction is not limited to H2B.8 as imbibition also induces the turnover of the H3.3 histone variant, thereby initiating a broad, replication-independent chromatin reprogramming process. These findings highlight a fundamental mechanism of epigenetic regulation during early plant development.

## Introduction

Seeds were a major innovation of photosynthetic organisms, contributing to the reproductive success of angiosperms. As they mature, seeds accumulate nutrients and reserves and acquire tolerance to desiccation, enabling long-term survival at low moisture levels. Despite its largely quiescent state, during which little transcriptional activity can be observed ^1,2^ the seed remains able to sense its environment and time germination to coincide with favorable environmental conditions, such as temperature and soil moisture content that are compatible with the completion of the plant’s life cycle. Reserve proteins and mRNA molecules accumulate during seed maturation, serving as templates for *de novo* protein synthesis ^3^. Protein biosynthesis from these stored mRNAs resumes upon water uptake during imbibition of the seed, as well as various metabolic and physiological activities, such as the mobilization of reserves to restart respiration and energy production, and the initiation of repair mechanisms ^4,5^. Imbibition also triggers changes in nuclear organization, as the volume of the small nuclei with highly condensed chromatin in the dry seeds of *Arabidopsis thaliana* progressively increases during water uptake and germination ^6^. The seed-to-seedling transition further requires transcription of new mRNAs, and more than 20,000 genes show differential expression ^7–9^, revealing important transcriptome reprogramming during this developmental transition.

Gene expression as a readout of the eukaryotic genome is strongly influenced by its chromatin organization, however, to date the contribution of local or higher-order chromatin organization to the observed transcriptome reprogramming during this developmental transition is only partly understood. At the level of the nucleosome, the basic building blocks of chromatin, post-translational modifications (PTMs) of histone proteins signal repressive or permissive chromatin states or directly influence nucleosome stability. The incorporation of specific histone variants has been demonstrated to further modify nucleosome stability or DNA accessibility and to influence the setting of histone PTMs ^10–12^. The Arabidopsis genome codes for multiple variants of H3, H2A, H2B core histones and the linker histone H1. This permits important combinatorial complexity, defining nucleosomes with different characteristics ^13,14^. The deposition, maintenance or exchange of histone variants is coordinated by a network of histone chaperones, which may exhibit specificity for a histone type and a particular variant. Histone chaperones such as FACILTATES CHROMATIN TRANSCRIPTION (FACT) ensure recycling of existing parental histones at the replication fork or during transcription ^15,16^, thereby permitting the retention of existing histone PTMs. Others such as CHROMATIN ASSEMBLY FACTOR 1 (CAF-1) and HISTONE REGULATOR A (HIRA) have been shown to coordinate *de novo* nucleosome assembly, thereby contributing to the reestablishment of specific histone modifications on the newly assembled nucleosome chains ^12^ or to maintain nucleosomal occupancy genome wide, respectively ^17–19^. The genomic distribution of histone variants has been investigated extensively in seedlings, revealing a correlation with specific chromatin states. For example, in seedling tissue, the replacement histone variant H3.3 is enriched at the 3’ end of transcriptionally active genes, while H2A.W marks transposable elements ^20–23^. During development, certain cell types express a particular histone variant repertoire, such as the small and highly condensed pollen sperm nuclei, which are enriched in H3.10 and the angiosperm-specific H2B variant H2B.8 ^24,25^. While both histone variants are not essential for male fertility, closer investigations showed that they fulfill specific functions. Due to amino-acid variations around the Lysine 27 site, H3.10 is a poor substrate for histone methyltransferases, and its incorporation in sperm chromatin contributes to reprogramming of the repressive H3K27me3 mark in paternal chromatin ^24^. H2B.8, in turn, is involved in nuclear and chromatin condensation of sperm nuclei, mainly through its phase separation capacity mediated by its long and intrinsically disordered N-terminal tail but does not influence the sperm cell transcriptome ^25^. Besides in sperm nuclei, the H2B.8 variant is expressed during seed maturation ^22,26^ during which chromatin condenses, and nuclear size progressively decreases^6^. Throughout this developmental time window, as DNA replication has ceased and in absence of *H3.1* expression, H2B.8 accumulates concomitantly with the replacement variant H3.3 ^27,28^. In dry seed, H3.3 is also enriched at 5’ ends of developmentally regulated genes and is required for their full activation during germination ^27^. Consequently, depletion of H3.3 in *H3.3* knockouts or reduced H3.3 incorporation in *HIRA* mutants leads to defects in germination and seed vigor ^27,28^. This led to the hypothesis that the seed-specific chromatin organization plays a role in genome protection and maintenance of the quiescent state, further suggesting that during germination and seedling establishment, changes in local and higher-order chromatin organization are required to achieve full transcriptome reprogramming.

Here, we have investigated the dry seed chromatin organization and determined the genomic distribution of the H2B histone variant H2B.8. We find that H2B.8 forms heterotypic nucleosomes at subset of genes and transposable elements on chromosome arms but is depleted from pericentromeric heterochromatin. The H2B.8-marked genes are globally repressed in dry seeds and co-enriched with gene-body H2A.Z and H3K27me3. While loss of H2B.8 does not affect the stored mRNA in dry seeds, during imbibition, the expression of a subset of H2B.8-marked genes is reduced in absence of H2B.8. 3D microscopy coupled to Hi-C analysis reveal that H2B.8 contributes to genome organization in the dry seed embryos, with H2B.8-enriched chromatin regions clustering in a H2B.8-dependent manner. Imbibition triggers degradation of stored *H2B.8* transcripts, and eviction of this histone H2B variant in a process that is independent of DNA replication but requires translation and *de novo* transcription. We find that histone eviction is not restricted to H2B.8, revealing global replication-independent chromatin reprogramming during the early stages of seed germination.

## Results

### H2B.8 occupies silent genes marked by H3K27me3 and H2A.Z

To study the distribution of the H2B.8 histone variant in dry seeds, we generated transgenic plants that express H2B.8-GFP under its endogenous promoter. Like H3.3-GFP ^29^, H2B.8 is strongly enriched in the small nuclei of the dry seed embryos (*Figure 1a*). In both cotyledons and hypocotyls, H2B.8 is present throughout the nucleus and in intensively stained chromatin domains (*Figure 1b*). Given the size and localization of some of these H2B.8-enriched domains at the nuclear periphery, these may coincide with chromocenters that comprise pericentromeric heterochromatin and nucleolus organizer regions (NORs). To investigate this and ensure that the genomic distribution of H2B.8 was not affected by the epitope tag position, we generated additional transgenic lines, in which the variant is fused either to a N-terminal Myc tag or to a C-terminal Flag tag. Simultaneous detection by immunostaining of the H2B.8 and the H2A.W histone variants, the latter marking pericentromeric heterochromatin ^21^, in isolated nuclei from dry seed embryos, revealed that H2B.8 is excluded from the H2A.W-enriched domains (*Figure 1c, Figure S1*). In dry seeds, H2B.8 is therefore enriched in larger euchromatic domains, while being globally depleted from pericentromeric heterochromatin.

**Figure 1:**
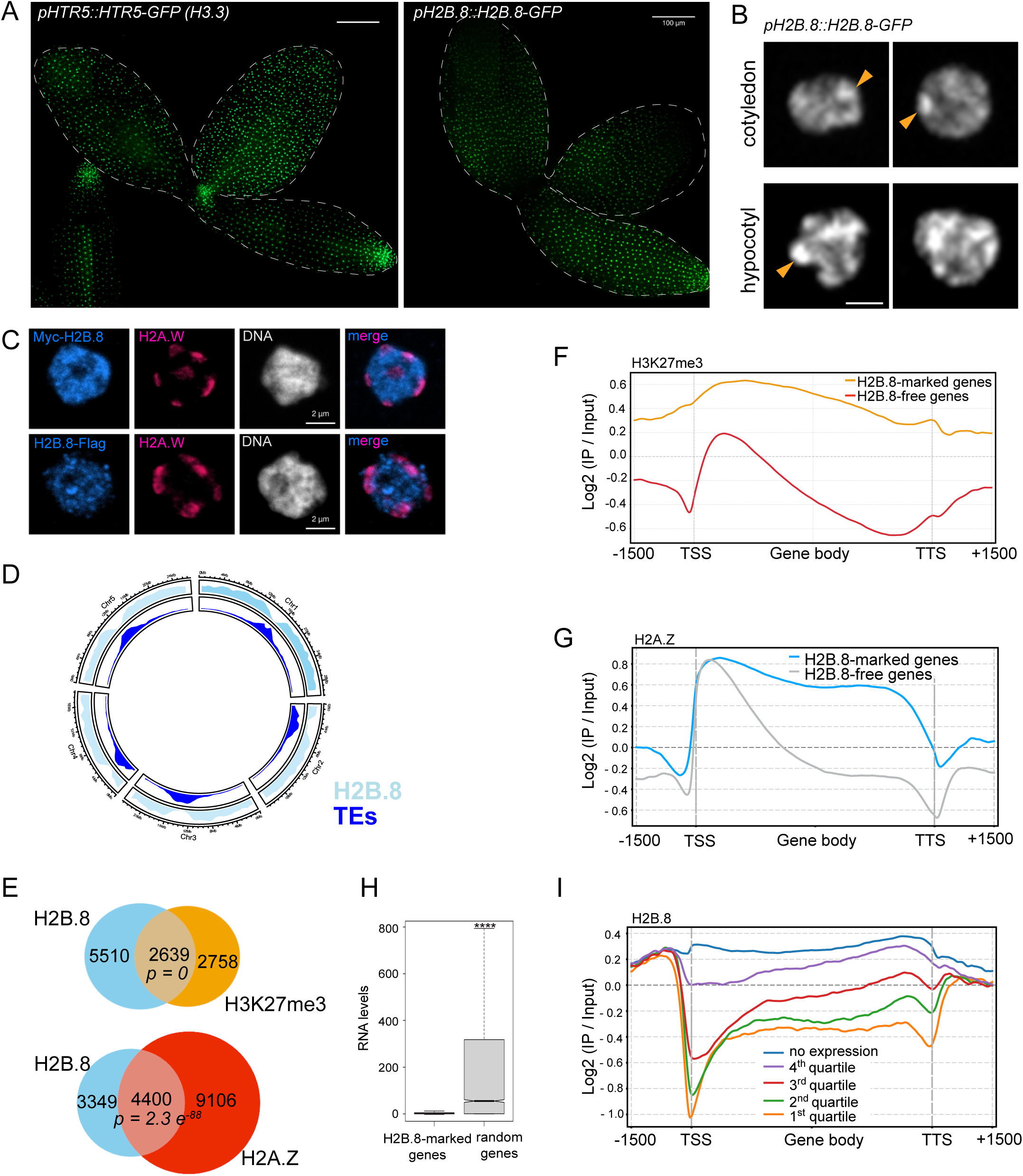
H2B.8 marks silent genes enriched in H3K27me3 and gene-body H2A.Z. **A.** Maximum intensity projections of dissected dry seed (DS) embryos expressing H3.3 (HTR5) or H2B.8 as a GFP fusion protein under the respective endogenous promoters. Embryos were dissected from the seed coat and imaged with a Spinning confocal microscope, 25x objective, and mosaics stitched together using the Zeiss software. **B.** Selected DS nuclei from the *pH2B.8::H2B.8-GFP* transgenic line imaged with a Spinning confocal microscope using a 63x objective. Arrowheads mark nuclear domains enriched in H2B.8-GFP both in cotyledons (top) and hypocotyl nuclei (bottom). **C.** Isolated DS embryo nuclei stained with DAPI (grey), in which H2A.W (magenta) and H2B.8 (blue) as a C-terminal Flag or N-terminal Myc fusion were revealed by immunostaining. **D.** Circos plot showing enrichment in H2B.8 (sky-blue) along the 5 chromosomes and relative to the centromeric / pericentromeric regions enriched in transposable elements (dark blue). **E.** Venn diagrams revealing a significant overlap between H2B.8-marked genes and genes enriched in H3K27me3 or H2A.Z in DS; p-values were determined with a hypergeometric test. **F.** Metagene plot of H3K27me3 distribution over all H2B.8-free and H2B.8-marked genes (n=7549). **G.** Metagene plot of H2A.Z distribution over all H2B.8-free and H2B.8-marked genes (n=7549). **H.** mRNA levels (DESeq2 normalization) of H2B.8-enriched genes and an equivalent number (n=7549) of random H2B.8-free genes, determined by RNA-seq in DS, *\*\*\*\* p<0.0001*, Mann Whitney test. **I.** Metagene plot showing profiles of H2B.8 enrichment over silent (absence of reads) and genes with different expression levels in DS.

To determine the genomic localization of this histone variant in dry seeds in more detail, we carried out ChIP-seq using a transgenic line expressing Myc-H2B.8 under control of its endogenous promoter. A set of common peaks was obtained from two independent replicates obtained with two different anti-Myc antibodies (*Figure S2a*). H2B.8 peaks occupy about 14.6% of the genome in dry seed embryos and, consistent with the immunofluorescence staining, H2B.8 is predominantly found along chromosome arms and is depleted from centromeric and pericentromeric heterochromatin (*Figure 1d*). Despite depletion of H2B.8 from the transposon (TE)-rich pericentromeric heterochromatin, 8713 TEs situated on chromosome arms show H2B.8 occupancy without preference for class I or class II TEs (*Figure S2b*). Besides TEs, we identified 7749 genes with H2B.8 peaks. 40% and 33% of these H2B.8-marked genes are respectively shared among H2B.8 targets in seedlings, which ectopically express H2B.8 ^25^, and sperm cells while the others are seed-specific (*Figure S2c, d*). To understand whether H2B.8 colocalizes with other histone variants or specific histone modifications, we retrieved available ChIP-seq data from dry seeds ^27,30^. Comparison of the ChIP-seq profiles showed that H2B.8 is globally depleted from genes carrying H3.3 or H3K4me3 peaks (*Figure S2e*). Instead, ∼49% of the genes carrying the repressive Polycomb modification H3K27me3 are marked by H2B.8 (*Figure 1e, f*); and ∼57% of all H2B.8-marked genes show H2A.Z enrichment (*Figure 1e*). This partial co-localization of H2B.8 with H2A.Z or H3K27me3 is also observed by immunofluorescence staining in dry seed embryo nuclei (*Figure S2f*). Closer investigation of the H2A.Z distribution reveals that H2B.8-marked genes show H2A.Z enrichment in the gene body (*Figure 1g*), a pattern that is associated with transcriptional repression ^31,32^. In addition, H2B.8 is depleted from accessible regions determined by ATAC-seq ^9^ (*Figure S2g*).

Given the co-enrichment of H2B.8-marked genes with H3K27me3 and gene-body H2A.Z, we expected H2B.8 to label genes that were transcriptionally repressed during seed maturation. To ascertain this, we analyzed the stored mRNA pool of wild type (WT) dry seeds. Comparison of mRNAs from genes occupied by H2B.8 to a random set of genes, revealed that for H2B.8-marked genes little to no stored mRNA is present in dry seeds (*Figure 1h*). Consistently, metagene analyses of H2B.8 enrichment as a function of transcript levels confirms the depletion of H2B.8 from genes, for which mRNAs accumulated during seed maturation (*Figure 1i, Figure S2h*). Taken together, H2B.8 marks TEs on chromosome arms and repressed genes enriched in H3K27me3 and/or gene-body H2A.Z.

### H2B.8 forms heterotypic nucleosomes

To characterize endogenous H2B.8, we generated a polyclonal antibody against a peptide in the N-terminal tail specific to H2B.8 (*Figure S3a*). By Western Blot analysis, this antibody detects the endogenous protein in WT plants as well as the Myc-tagged version of H2B.8 in the transgenic line (*Figure 2a*). We noticed that both the endogenous as well as the tagged H2B.8 migrate at a considerable higher molecular weight than the predicted 27kDa for the endogenous protein. To test whether the unexpected electrophoretic mobility is caused by post-translational modifications set *in planta* ^33^, we expressed full-length H2B.8 or only its N-terminal tail as Gal4-DNA binding domain fusions with a Myc-tag in yeast. Both fusion proteins migrate at a similar molecular weight and slower compared to the H2B.4 variant (*Figure S3b*), revealing that, in dry seeds, the reduced electrophoretic mobility of H2B.8 is conferred by the tail sequence rather than by post-translational modifications.

**Figure 2:**
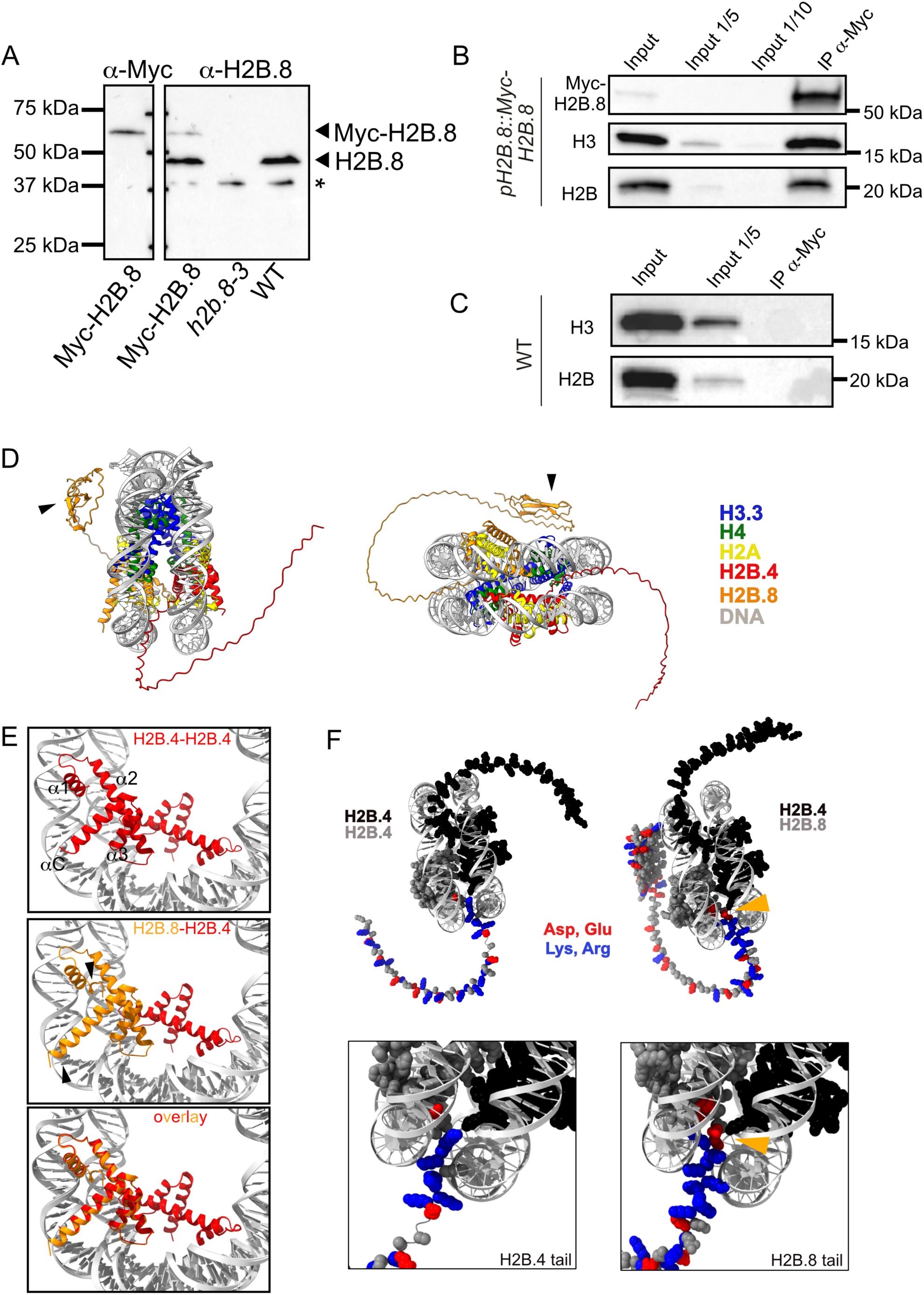
H2B.8 forms heterotypic nucleosomes. **A.** Western Blot using an *in house* generated anti-H2B.8 antibody. From left to right: Nuclear extracts from seeds expressing 4xMyc-H2B.8 under its endogenous promoter in the WT background, *h2b.8-3* mutant and WT seeds. The anti-Myc antibody (left) detects Myc-tagged H2B.8. Myc-tagged H2B.8 and endogenous H2B.8, revealed by the anti-H2B.8 antibody, are indicated with arrowheads. * = non-specific band. **B-C.** Immunoprecipitation of nucleosomes with an anti-Myc antibody from dry seeds expressing 4xMyc-H2B.8 (**B**) or WT dry seeds (**C**). Revelation with an anti-Myc, anti-H3 and anti-H2B antibody reveals the presence of H3 and smaller H2B variants in the Myc-H2B.8 containing nucleosomes. **D**. Structure of the heterotypic nucleosome predicted by AlphaFold 3 comprising H3.3 (blue), H4 (green), H2A.1 (yellow), H2B.4 (red) and H2B.8 (orange). Part of the H2B.8 N-terminal tail (AA8 - 65) is predicted to fold into β-sheets (arrowheads). The N-and C-terminal tails of all histones except H2B.4 and H2B.8 have been hidden for clarity. **E.** Sub-fraction of the predicted nucleosome structures with either two H2B.4 molecules (red, top) or one H2B.4 and one H2B.8 (red and orange, middle) and the overlay of the two structures (bottom). H2B.8 is predicted to form a longer α1 and an extended C-terminal helix (arrowheads). Only the alpha-helical, structured parts of the H2B histone variants are shown. **F.** Sub-fraction of the predicted nucleosome structures with either two H2B.4 molecules (left) or one H2B.4 and one H2B.8 (right) centered on the H2B N-terminal tail that protrudes between the two DNA gyres. Acidic amino acids (ASP, GLU) and basic amino acids (LYS, ARG) are colored in red and blue respectively. The arrowhead indicates additional acidic amino acids in the H2B.8 tail.

The incorporation of different histone variants into nucleosomes results in important combinatorial complexity ^13^. However, not all combinations are favorable as for example H2A variants form exclusively homotypic nucleosomes ^34^. To investigate whether H2B.8 nucleosomes in the dry seed are homotypic (two H2B.8 molecules per nucleosome) or heterotypic (one H2B.8 and another, shorter H2B variant), we immunoprecipitated nucleosomes from dry seeds expressing Myc-H2B.8. Mononucleosomes were released by MNase digestion (*Figure S3c*) and H2B.8-containing nucleosomes immunoprecipitated with an anti-Myc antibody. Histone H3 was detected only in the immunoprecipitated fraction from seeds expressing Myc-H2B.8, but not from WT seeds, showing that nucleosomes were successfully and specifically precipitated (*Figure 2b, c*). Small H2B variants were detected by a generic H2B antibody in a similar IP to input ratio as H3. Thus, H2B.8 forms heterotypic nucleosomes, in which H2B.8 is associated with one of the shorter H2B variants.

Based on this observation, we predicted nucleosome structures with AlphaFold 3 comprising a 146bp long Widom 601 sequence, H3.3, H4, H2A.1, H2B.4, and H2B.8 (*Figure 2d*). We chose H3.3, H2A.1 and H2B.4 as these variants are highly expressed in seeds (^27,28^ and *Figure S3d & S5a*). AlphaFold predicts, although with lower confidence, the presence of a small, structured domain with β-sheets in the very N-terminal region of H2B.8 (AA8-65, *Figure 2d*) within the otherwise intrinsically disordered N-terminal tail (AA66-145). Other differences of H2B.8 compared to the other 10 H2B variants include changes from basic to acidic amino acids within the second and third alpha helices (*Figure S3a*), which form contacts in the H2A-H2B dimer and the H4-H2B four-helix bundle ^35^ and the presence of an extended C-terminal tail. We therefore analyzed the histone fold domain in the predicted nucleosomes comprising either H2B.4 and H2B.8 or two H2B.4 molecules. Compared to H2B.4, H2B.8 forms a longer structured α1 helix as well as an extended C-terminal alpha helix (*Figure 2e*). Finally, just N-terminally of the first alpha helix and before a longer stretch of basic Lysines and Arginines, H2B.8 comprises an insertion of 5 amino acids including two acidic amino acids (*Figure S3a*), which are situated in close contact to the two DNA gyres (*Figure 2f*).

Thus H2B.8 in dry seeds forms predominantly heterotypic nucleosomes with one of the shorter H2B variants. In these heterotypic nucleosomes, H2B.8-specific features are predicted to lead to distinct histone-histone or histone-DNA interactions in the H2B.8-comprising nucleosome.

### H2B.8 influences 3D chromatin organization in dry seeds

To understand whether H2B.8 affects the transcriptome and shapes the organization of chromatin in dry seeds, we obtained a T-DNA insertion mutant, *h2b.8-3*, in which the T-DNA is situated in the exon, and which lacks the H2B.8 protein (*Figure 2a*). After-ripened seeds of *h2b.8* mutants germinate with comparable kinetics to WT under light (*Figure S4a*) or when stimulated by Gibberellin or Ethylene in the dark (*Figure S4b, c*) and H2B.8 is dispensable for the increased dormancy observed upon loss of the histone chaperone HIRA ^28^ (*Figure S4d*). To determine whether H2B.8 deposition affects the stored mRNA pool in dry seeds, we analyzed the transcriptome by RNA-seq in single *h2b.8* mutants as well as in the *hira-1* mutant background, in which histone H3 occupancy is reduced ^28^. Transcript levels of 991 genes are modified in absence of HIRA (*Figure 3a*), with 334 showing reduced and 647 increased transcript levels in the *hira-1* mutant, among the latter numerous histone genes (*Figure S5a*). Deficient nucleosome assembly therefore affects transcriptional regulation during seed maturation and consequently the stored mRNA pool. Instead, RNA levels of genes, including H2B.8-marked genes, are unaffected by loss of H2B.8 (*Figure 3a, Figure S5b*), and the *hira-1 h2b.8-3* double mutant transcriptome strongly resembles the one of *hira-1* single mutants consistent with the observations in sperm nuclei ^25^ and the seed germination phenotype (*Figure 3a, Figure S5c, d, Figure S4d*). Therefore, while deficient H3.3 deposition alters the dry seed transcriptome, H2B.8 deposition is dispensable for the establishment of the mRNA pool in dry seeds ^25^.

**Figure 3:**
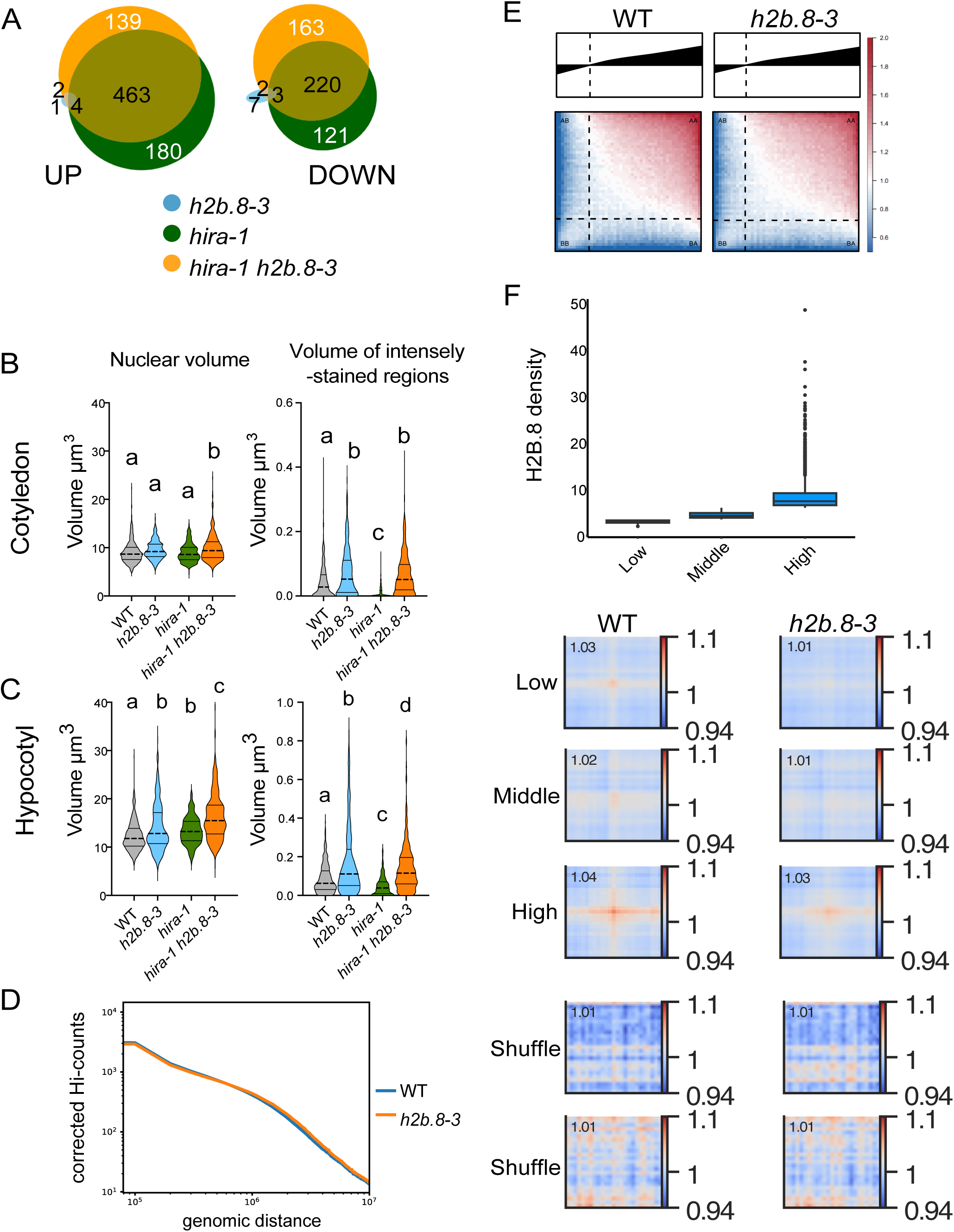
H2B.8 regulates 3D chromatin organization in dry seeds. **A**. Venn diagram showing up-and downregulated genes (*log2FC > 1, padj < 0,05*) in DS of *h2b.8-3*, *hira-1* and the *h2b8-3 hira-1* double mutants relative to WT. **B**, **C**. Nuclear volume and total volume of intensely stained chromatin regions per nucleus in cotyledon (**B**) and hypocotyl nuclei (**C**) from dissected DS. Nuclear parameters were determined using a deep-learning model generated to predict nuclear shape and Hoechst-dense regions using the *biom3d* deep learning framework. Cotyledons: WT: n (number of nuclei) = 420, N (number of embryos) = 21, *h2b.8-3*: n= 260, N =13, *hira-1*: n= 300, N= 15 and *hira-1 h2b.8-3*: n= 500, N= 25. Hypocotyls: WT: n = 440, N = 22, *h2b.8-3*: n= 300, N =15; *hira-1*: n= 340, N= 17 and *hira-1 h2b.8-3*: n= 415, N= 21. Significant differences (p<0.05) between genotypes were determined using an ordinary one-way ANOVA coupled to a Tukey’s multiple comparisons test. **D**. Averaged scaling plots with a 100 kb genomic bin size, illustrating how interaction frequencies vary with increasing genomic distance across all chromosomes. **E**. Saddle plots showing chromatin compartmentalization in DS from WT and *h2b.8-3*. The mean cis observed/expected interaction frequencies between 20 kb bins are plotted, ordered by PC1 values from WT. **F**. Top: Boxplot of H2B.8 density showing three quantiles of H2B.8 peaks, classified as high, middle, and low H2B.8 enrichment based on fold change relative to input. Bottom: Aggregate plots showing the mean contact matrices for WT and *h2b.8-3* at peaks grouped into three H2B.8 enrichment quantiles. Contacts in *h2b.8-3* are noticeably lower compared to WT. The two shuffle rows represent randomly selected genomic regions, serving as background controls.

Given that H2B.8 contributes to chromatin condensation in sperm nuclei ^25^, we then sought to investigate whether H2B.8 incorporation shapes the nuclear morphology and influences higher-order chromatin organization in dry seed embryos. To quantify nuclear morphology and the condensed chromatin regions that intensely stain with DNA dyes, 3D images of cotyledon and hypocotyl nuclei were obtained from Hoechst-stained dry seed embryos of WT, *h2b.8-3, hira-1* and *h2b.8-3 hira-1* double mutants. A deep-learning model was trained on a set of manually annotated images from embryos to segment nuclei and intensely-stained chromatin regions (*Figure S6a*). While the shape of cotyledon and hypocotyl nuclei remains unaltered in absence of HIRA and/or H2B.8 (*Figure S6b, c*), we observed a modest increase in the volume of hypocotyl nuclei in *h2b.8* and *hira-1* mutants, as well as an additive effect on nuclear volume in both cotyledons and hypocotyls of seeds lacking both HIRA and H2B.8 (*Figure 3b, c; Figure S6c*). Next, we segmented the intensely stained chromatin regions and determined their volume per nucleus. While fewer chromatin aggregates are detectable in *hira-1* mutants, loss of H2B.8 both in the WT and the *hira-1* mutant background results in more discernable heterochromatin regions in both cotyledon and hypocotyl nuclei compared to WT (*Figure 3b, c; Figure S6c*). H2B.8 therefore moderately contributes to nuclear size, and its absence leads to altered chromatin organization in dry seeds. This effect is independent of global chromatin accessibility as determined by MNase digestion (*Figure S7*) and dominant over the microscopic changes in chromatin organization induced by loss of HIRA.

Next, to investigate the role of H2B.8 in organizing chromatin in dry seed nuclei in molecular detail, we generated high-resolution maps of 3D chromatin architecture in Arabidopsis by conducting Hi-C experiments from manually dissected WT and *h2b.8-3* mutant dry seed embryos (see *Supplementary Data 1* and Materials and Methods). Genome-wide analyses of eigenvalues, as well as correlation analyses of the first principal component (PC1) values (*Figure S8a*) together with the visual inspection of Hi-C interaction maps (*Figure S8b, c*) confirmed strong consistency between experiments. Using cLoops2 ^36^, we estimated the Hi-C resolution to 5 kb for both control samples and the *h2b.8-3* mutant (*Figure S9*). To evaluate whether loss of H2B.8 influences the global higher-order chromatin organization, we computed differential interaction maps, which showed no substantial changes in contact frequency in the mutant compared to the WT dry seeds (*Figure S10*). As chromatin interaction frequencies typically decrease with genomic distance according to a power-law function ^37^, we generated scaling plots for both genotypes, which revealed no detectable differences in the decay of interactions over distance, indicating that both short-and long-range contacts remain largely unaltered upon loss of H2B.8 (*Figure 3d*). Furthermore, comparison of PC1 profiles between WT and mutant samples showed no marked shifts between A (transcriptionally active) and B (inactive) compartment identities (*Figure 3e*). Thus, global chromatin compartmentalization in dry seed embryos appears largely stable in the absence of functional H2B.8. Previous work in both plant and animal systems suggests that chromatin regions marked by similar histone modifications or bound by common regulatory factors preferentially associate in 3D space ^38^ and H2B.8 has previously been shown to be involved in chromatin condensation of sperm nuclei ^25^. To determine whether H2B.8 contributes to such homotypic interactions in dry seed embryos, we stratified H2B.8-enriched regions in WT into three categories (low, medium, high) based on ChIP-seq signal intensity. We then constructed aggregate Hi-C submatrices to quantify mean contact enrichment between similarly marked regions. Following observed versus expected normalization, WT samples showed enriched contacts among H2B.8-marked loci, whereas this enrichment was reduced in the *h2b.8* mutant (*Figure 3f*). Together with the microscopic analysis of dry seed nuclei, these findings suggest that H2B.8 plays a role in facilitating spatial proximity among specific euchromatin domains.

### H2B.8 is rapidly evicted upon imbibition and impacts transcriptional reprogramming

Given that H2B.8 is enriched in embryo chromatin in dry seeds and contributes to its higher-order organization, our subsequent objective was to investigate the dynamics of this variant during seed germination. To this end, the expression and localization of H2B.8 was monitored in our experimental setup, which involved imbibing seeds for 48 hours at 4°C in the dark, followed by germination in a growth chamber set to long-day conditions. In this experimental setup the first radicle protrusion is observed around 48h after transfer to light (*Figure S4a*). During imbibition, as soon as 24h contact with water, *H2B.8* transcript levels strongly decrease (*Figure 4a; Figure S11a*). The observed, progressive downregulation of the *H2B.8* gene correlates with the transition from an H3K4me3-rich chromatin state in seeds to an H3K27me3-marked chromatin state in seedlings (*Figure 4b*). To characterize the *H2B.8* promoter activity in more detail, we expressed the *β-glucuronidase (GUS*) reporter gene from the *H2B.4* (control) or the *H2B.8* promoter and followed GUS activity using X-Gluc histochemical staining. The control construct driven by the *H2B.4* promoter shows that this promoter was active during maturation and results in strong *GUS* expression throughout germination (*Figure 4c; Figure S11a*). In contrast, the *GUS* gene driven by the *H2B.8* promoter is progressively downregulated during seed imbibition, with the loss of blue staining initiating in the cotyledons. After 24h in the light, only faint staining remains visible in the radicle (*Figure 4c*). Next, to investigate the fate of the H2B.8 protein during the seed-to-seedling transition, we carried out Western Blot analyses. Both Myc-tagged H2B.8 (*Figure S11b*) and endogenous protein levels (*Figure 4d*) decrease during imbibition in the cold with up to 50% of the H2B.8 variant protein being evicted and degraded after 24h. This contrasts with the long half-life of several weeks of mammalian histones in non-proliferating cells ^39,40^. To confirm these results by a different method, we investigated H2B.8 dynamics during imbibition by live cell imaging. In dry seeds, GFP-tagged H2B.8 can be detected in cotyledons, hypocotyls and strongly in the radicle. At 24h imbibition, H2B.8 has been evicted mainly in the cotyledons, at 48h faint H2B.8-GFP signals remain detectable in the root tip, and after 24h in the light, H2B.8-GFP is undetectable (*Figure 4e*). Therefore, consistent with the GUS staining that monitors *H2B.8* promoter activity, H2B.8 eviction takes place most rapidly in the cotyledons. Given the substantial co-enrichment in H2B.8 and H3K27me3 in dry seeds, we then asked whether H2B.8 occupancy or its subsequent eviction during imbibition influences H3K27me3 reprogramming during the seed-to-seedling transition, however, no significant overlap between the genes that gain or lose H3K27me3 ^41^, and H2B.8 could be detected (*Figure S12a*).

**Figure 4:**
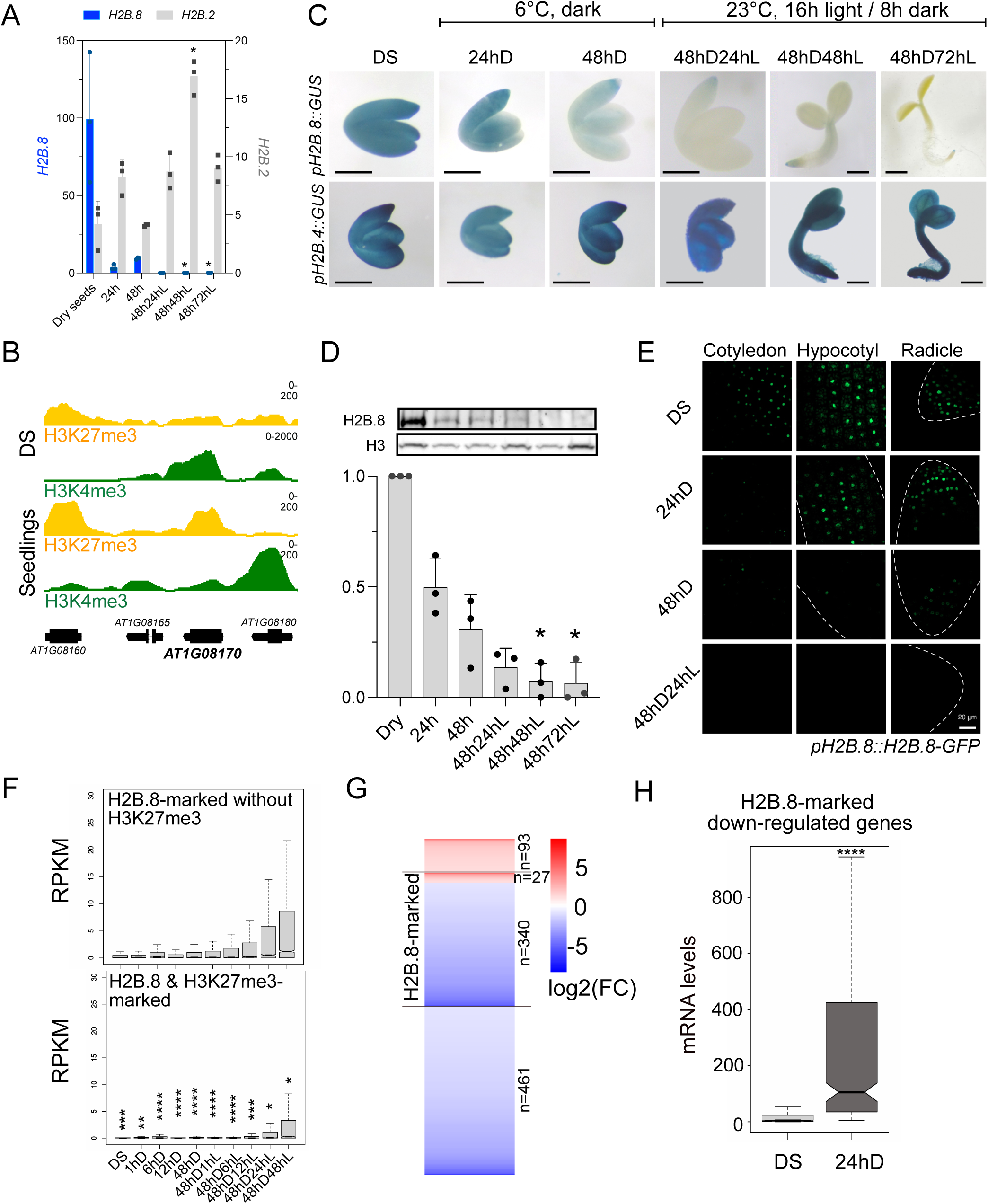
Seed imbibition triggers H2B.8 eviction. **A**. Relative transcript levels of *H2B.8* and *H2B.2* normalized to *MON1* and *TIP41* determined by RT-qPCR in DS, after 24h or 48h imbibition in the dark (24hD, 48hD) and in stratified seeds after 24h, 48h or 72h of incubation under long days (48hD24hL - 48hD72hL). *\* p<0.05*, Friedman’s test. **B**. Genome browser view of H3K4me3 and H3K27me3 at the *H2B.8* locus in dry seeds and 8-day old seedlings (ChIP-seq data were retrieved from ^30,74^). **C**. GUS-stained embryos in transgenic lines expressing *GUS* under control of the *H2B.8* (top) or the *H2B.4* (bottom) promoter. Time points as in **A**. Scale bar represents 0.5mm. **D**. Representative Western Blots of H2B.8 and the H3 loading control (top) and quantification (bottom) of endogenous H2B.8 normalized to H3 during the seed-to-seedling transition in four independent biological replicates at the same time points as in **A**. H2B.8 levels in dry seeds were set to 1. *\* p<0.05,* Friedman’s test coupled to a Dunn’s multiple comparisons test. **E**. Representative maximum intensity projections of cotyledons, hypocotyls or radicles from embryos expressing H2B.8 as a GFP fusion under control of its endogenous promoter. Embryos were dissected from DS, imbibed and germinating seeds at 24hD, 48hD and 48hD24hL and imaged with a Spinning confocal microscope, 63x objective. Scale bar represents 20 μm. **F**. Transcript levels in RPKM during the seed-to-seedling transition (RNA-seq dataset from ^8^) of genes marked by H2B.8 but not H3K27me3 (n=4956) or both H2B.8 and H3K27me3 (n=2592, H3K27me3 ChIP-seq dataset from ^30^) in DS. *\* p< 0.1, **p< 0.05, *** p<0.01, **** p<0.0001*, non-parametric Mann-Whitney test. **G**. Heat map showing up and down-regulated genes after *spike-in* normalization in *h2b.8-3* mutants compared to WT at 24hD. 42% (340 / 801) of the down-regulated genes are marked by H2B.8. **H**. Transcript levels of H2B.8-marked genes (n=340), which are down-regulated in *h2b.8-3* mutants at 24hD, in WT DS and at 24hD imbibed seeds, *\*\*\*\* p< 0.0001*, Wilcoxon matched-pairs signed rank test.

To study the global transcriptional dynamics of H2B.8-marked genes, we determined their expression pattern during the seed-to-seedling transition in an available RNA-seq dataset ^8^. H2B.8-marked genes are globally expressed to low levels during germination and seedling establishment (*Figure S12b*), however, compared to genes marked by both H2B.8 and H3K27me3, a subset of genes marked only by H2B.8 is progressively activated during seed germination (*Figure 4f*). To understand whether the presence of H2B.8 in dry seeds or its dynamics during imbibition affect transcriptional reprogramming, we carried out *spike-in* RNA-seq analysis of WT and *h2b.8-3* mutant seeds harvested at 24hD. We find that 921 genes are differentially regulated in imbibed *h2b.8-3* mutant seeds (*Figure S13c*) with the majority (801) being downregulated compared to WT. More than 40% (340) are marked by H2B.8 in dry seeds (*Figure 4g*) and these genes are globally induced during imbibition (*Figure 4f*). Together this indicates that the transcriptional activation of a subset of H2B.8-marked genes is influenced by the enrichment in this histone variant, either by pre-marking these genes for activation or by facilitating their activation through H2B.8 eviction. Therefore, concomitant with transcriptional reprogramming, seed imbibition triggers degradation of *H2B.8* transcripts and eviction of H2B.8 histone variants from chromatin revealing that water uptake initiates chromatin reprogramming.

### Histone eviction during imbibition is not restricted to H2B.8

Given the deposition and eviction patterns of H2B.8 histones, their assembly into nucleosomes may involve specific histone chaperone machineries that are expressed specifically during seed maturation. To understand whether this is the case, we expressed Myc-H2B.8 under the control of the *HTR5 (H3.3)* promoter and monitored H2B.8 protein levels during the seed-to-seedling transition. While H2B.8 is initially evicted, H2B.8 can, when continuously expressed, be re-deposited during germination (*Figure 5a*), demonstrating that its incorporation is mediated by ubiquitous histone chaperone systems. One of the histone chaperones that handle H2A-H2B histones is FACILITATES CHROMATIN TRANSCRIPTION (FACT) ^42^. To determine whether FACT plays a role in H2B.8 dynamics, we determined H2B.8 levels in WT and *ssrp1-2* mutants, which express a truncated SSRP1 subunit of the FACT complex ^43,44^. In dry seeds, relative H2B.8 protein levels normalized to H3 are reduced in *ssrp1-2*, but not in *hira-1* mutants compared to WT (*Figure 5b*), revealing that FACT is important for H2B.8 deposition or maintenance in the dry seed.

**Figure 5:**
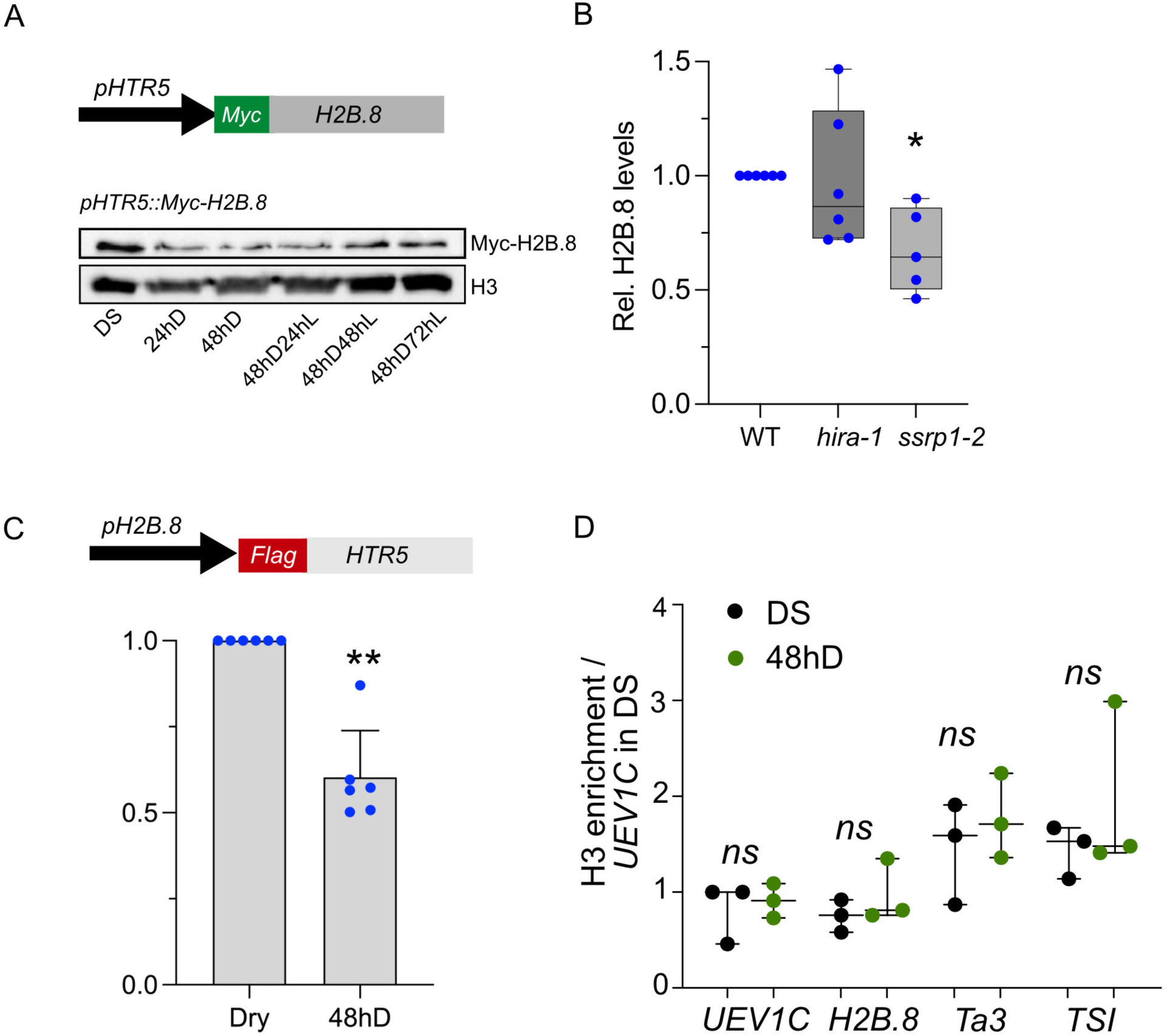
Histone eviction is not restricted to H2B.8. **A**. Representative Western Blot showing expression of Myc-tagged H2B.8 under control of the *HTR5 (H3.3)* promoter during the seed-to-seedling transition together with the H3 loading control. **B**. Quantification of endogenous nuclear H2B.8 proteins relative to H3 in WT, *hira-1* and *ssrp1-2* mutant seeds. Graph shows mean of ≥ 5 biological replicates. *\* p= 0.0537*, non-parametric Friedman test coupled to a Dunn’s multiple comparisons test. **C**. Quantification, relative to total H3, of endogenous H2B.8 and Flag-H3.3 expressed under the *H2B.8* promoter in DS and after 48h imbibition in the dark (48hD). Graph shows mean of 5 independent transgenic lines. *\*\* p< 0.05,* non-parametric Wilcoxon test. Error bars correspond to SD. **D**. H3 ChIP-qPCR analysis (n = 3) in DS and seeds stratified for 48h in the dark (48hD). Values were normalized to input and to the H3 levels at the *UEVC (At2g36060)* gene in DS, *ns = not significant*, ttest.

Considering the observed eviction and degradation of H2B.8 during imbibition, we asked whether this eviction is representative of global chromatin reprogramming or whether the observed histone eviction is restricted to H2B.8. To this end, transgenic lines were established, which express Flag-H3.3 under control of the *H2B.8* promoter. We expect that in these lines, following the immediate downregulation of the *H2B.8* promoter (*Figure 4a, c*) little to no new Flag-H3.3 will be produced upon imbibition. In consequence, the dynamics of the *’seed’* H3.3 pool, deposited during seed maturation, can be followed. We find that, in multiple independent transgenic lines, Flag-H3.3 levels were decreased 48 hours after imbibition in comparison to the dry seed (*Figure 5c*) indicating that eviction of *’seed’* histones during imbibition is not exclusive to H2B.8. To investigate whether eviction of the *’seed’* H3.3 histones is compensated by *de novo* histone deposition during imbibition, we conducted H3-ChIP-qPCR to evaluate nucleosome occupancy at 48hD. At the analyzed loci, there is no evidence of reduced nucleosome occupancy at 48hD (*Figure 5d*), therefore eviction of *’seed’* H3.3 is compensated by deposition of new H3 histones. In conclusion, seed imbibition is characterized by significant histone eviction and *de novo* deposition, indicating substantial chromatin reprogramming during the initial stages of seed germination.

### Histone H2B.8 eviction takes place in a DNA-replication independent manner but requires de novo transcription and translation

The process of DNA replication presents an opportunity for modification of chromatin structure, as new histones are deposited concurrently with DNA replication. The progressive loss of *‘seed’* histones (*Figure 4d, e, Figure 5c*) may therefore be attributed to the dilution of the existing histones by the newly deposited histones during the process of DNA replication. To investigate whether the chromatin reprogramming that occurs during imbibition is replication-coupled, we timed the onset of DNA replication in embryos in our experimental setup. Flow cytometry analysis of dissected embryos at different time points during the seed-to-seedling transition revealed one sharp peak in dry seeds, showing that most embryonic nuclei are arrested in G1 phase (*Figure 6a*). The initial broadening of the peak and the emergence of S-phase nuclei at 24 hours after transfer to light indicate that DNA (endo)replication initiates between 48hD and 48hD24hL (*Figure 6a*) before a distinct G2 peak emerges by 48hD48hL. The onset of DNA replication was confirmed by EdU-staining, which revealed the first EdU-positive nuclei at 48hD24hL, predominantly in the radicle (*Figure 6b*). Therefore, the chromatin reprogramming leading to the eviction of H2B.8 and H3.3 during imbibition during the first 48h of imbibition (*Figure 4d, e; Figure 5c*) occurs independently of DNA replication.

**Figure 6:**
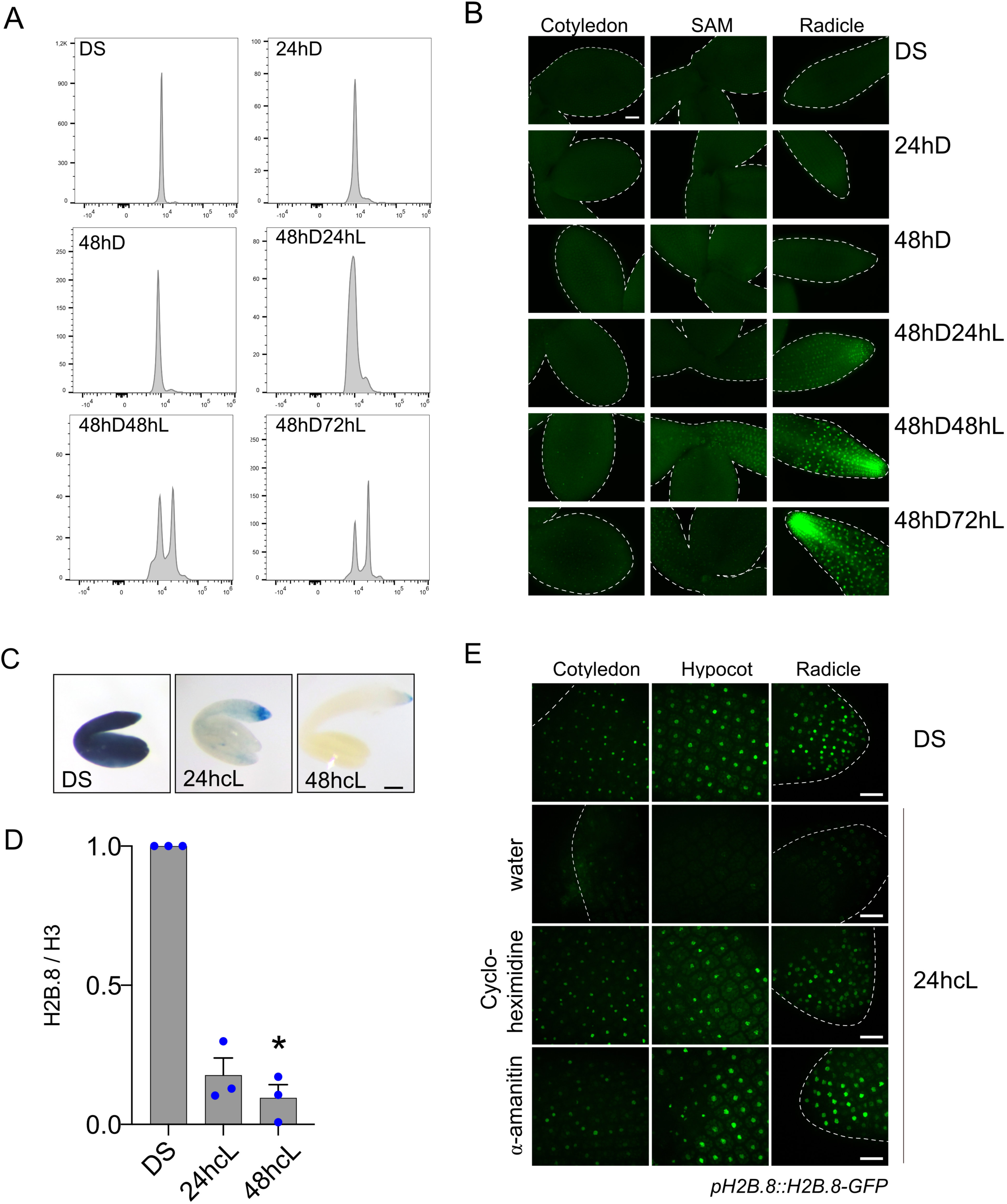
H2B.8 eviction during imbibition takes place in a DNA-replication independent manner but requires translation and transcription. **A**. Flow cytometry analysis of nuclei from isolated embryos at the indicated stages during the seed-to-seedling transition. **B**. EdU incorporation revealed by Click-IT chemistry in embryos at different stages during the seed-to-seedling transition. Images were acquired with a Spinning confocal microscope, equipped with a 63x objective. **C**. β-glucuronidase (GUS) histochemical assay revealing activity of the *H2B.8* promoter in transgenic lines expressing *GUS* under control of the *H2B.8* promoter in DS, and after 24h or 48h incubation under continuous light (24hcL, 48hcL). **D**. Quantification from Western Blots (n=3) of endogenous, nuclear H2B.8 normalized to H3 levels in DS, at 24hcL and 48hcL. *\* p<0.05,* Friedman’s test coupled to a Dunn’s multiple comparisons test. **E**. Representative maximum intensity projections of cotyledons, hypocotyls and radicles from embryos expressing H2B.8 as a GFP fusion under its endogenous promoter isolated from dry seeds or imbibed seeds under continuous light (24hcL). Images were acquired with a Spinning confocal microscope, 63x objective using identical exposure settings for treated and untreated samples. Seeds were imbibed in water or in the presence of 100μM Cycloheximidine or 500μM α-amanitin.

Based on the dynamics of H3.3 and H2B.8 eviction, we hypothesized that chromatin reprogramming reflects the physiological state of the seed. To test this, we exposed imbibed seeds under continuous light (cL). Under these conditions, 25 to 57% of the seeds germinated after 48h across the three biological replicates. GUS staining is diminished at 24hcL compared to 24hD (*Figure 6c, 4c)* revealing rapid *H2B.8* promoter inactivation and *GUS* transcript degradation. Western Blots further reveal that about 80% of H2B.8 proteins are evicted and degraded after 24hcL (*Figure 6d*). H2B.8 degradation at least partly involves the proteasome pathway, as H2B.8-GFP is retained longer in seeds treated with proteasome inhibitors (*Figure S13a*). The observed differences in H2B.8 eviction between imbibition in dark/cold and light/23°C provide evidence that chromatin reprogramming occurs concomitantly with the physiological alterations during the seed-to-seedling transition. Germination requires translation of stored mRNAs, while *de novo* transcription is not essential for radicle protrusion ^5^. To test whether H2B.8 eviction during imbibition necessitates the capacity of the embryo to translate stored mRNAs or to initiate *de novo* transcription, seeds expressing H2B.8-GFP were treated with the translational inhibitor cycloheximide or with alpha-amanitin that inhibits RNA Polymerase II activity. Both treatments impede H2B.8 eviction and degradation (*Figure 6e*), demonstrating that protein synthesis and the global transcriptional onset upon water uptake are necessary for the observed chromatin reprogramming in the embryo during imbibition. Therefore, seed imbibition triggers replication-independent chromatin reprogramming requiring protein translation and transcriptional activity.

## Discussion

### H2B.8 incorporation modifies chromatin organization in dry cell nuclei

Dry seed embryos in Arabidopsis present small nuclei with highly condensed chromatin ^6^. Here, we demonstrate that the embryonic nuclei in dry seeds are characterized by the presence of the H2B.8 histone variant, which accumulates in transcriptionally repressed chromatin regions that cluster in 3D nuclear space, and is evicted in a replication-independent manner upon water uptake during imbibition. Expressed late during seed maturation, H2B.8 accumulates in dry seeds at TEs located on chromosomal arms as well as at H3K27me3 and gene body H2A.Z-enriched genes suggesting the existence of locus specific deposition mechanisms. Given that H2B.8 is expressed during the maturation phase ^45^ by the time proliferation has ceased ^46^ and accumulates at silent loci, both replication-and transcription-coupled H2B.8 deposition are unlikely to be the predominant modes of H2B.8 chromatin assembly. Instead, the observed H2B.8 enrichment patterns in dry seeds could rather reflect its accumulation in chromosomal regions that are devoid of active transcription but undergo histone H2A-H2B exchange catalyzed by chromatin remodeling factors as observed for the deposition of H2A.Z that relies on the SWR1 chromatin remodeling complex ^47^ or H2A.W that requires DECREASE IN DNA METHYLATION1 ^48,49^. Reduced H2B.8 levels in a mutant for histone chaperone complex FACT, known to reorganize nucleosomes, by disrupting histone-histone and DNA-histone interactions and by depositing new or recycled histone H2A-H2B dimers ^50,51^ suggests a role for FACT in H2B.8 maintenance during seed maturation. Furthermore, the exclusion from the pericentromeric regions marked by H2A.W variants and the co-enrichment with H2A.Z may indicate that specific histone variants combinations within a nucleosome are favored.

H2B.8 differs from the other H2B variants by several sequence features, and its incorporation is therefore likely to affect nucleosome structural properties. Modified nucleosome properties due to H2B variant incorporation have been observed for different H2B variants in other organisms, e.g. upon deposition of the primate specific H2B.W1 variant, which is expressed in spermatogonia. Like H2B.8, H2B.W1 has an extended C-terminal tail, and it exhibits weakened histone-DNA interactions ^52^. Another example is H2BE, expressed in olfactory epithelium cells of the mouse brain ^53,54^, where it increases chromatin accessibility, a property conferred by a single amino-acid change (V39I) in the α1 helix at the interface between the histone octamer and DNA ^54^. At this position H2B.8 carries an arginine (I153R, *Supplementary Figure 13b*) that points towards the DNA suggesting stronger DNA-histone interactions. Future *in vitro* studies should help to better understand to what extent the incorporation of H2B.8 into the nucleosome affects its stability or DNA unwrapping properties in different heterotypic nucleosomes.

A common feature of sperm and dry seed nuclei that express H2B.8 is their small size ^6,28^, suggesting that H2B.8 could contribute to the tight chromatin organization in seeds through its phase separation properties ^25^. In dry seeds, loss of H2B.8 only moderately affects the volume of embryonic nuclei. Incorporation of H2B.8 is therefore not the major determinant of chromatin condensation during seed maturation and the changes in nuclear volume during germination might not be regulated by eviction of H2B.8, but rather by the nuclear lamina components CROWDED NUCLEI (CRWN), which are required for the increase in nuclear volume during germination ^6^. Instead, H2B.8 incorporation contributes to the higher-order organization of chromatin in embryonic nuclei. In seeds, large H2B.8-enriched chromatin domains can be visualized both by live-cell imaging and immunofluorescence. Our Hi-C analysis further revealed that H2B.8-rich domains tend to cluster within the 3D nuclear space. Although the global chromatin architecture is not markedly altered in the absence of H2B.8, the 3D clustering of H2B.8-enriched regions along chromosome arms is diminished in seeds lacking this histone variant. As a result, other higher-order chromatin domains such as the H2B.8-depleted pericentromeric regions may be more easily visualized by microscopy and segmented from the remaining condensed euchromatin.

As in sperm cells ^25^ and consistent with the co-enrichment in gene-body H2A.Z and H3K27me3, H2B.8 marks lowly expressed and silent genes in embryos, a genomic distribution, which is likely linked to its deposition mode as discussed above without however being required for the transcriptional repression of these genes. In seedlings, transcriptional activation correlates with eviction of ectopically expressed H2B.8 ^25^. We therefore hypothesized that H2B.8 eviction initiated upon water uptake in embryos could facilitate transcriptional reactivation during imbibition. This hypothesis is indeed supported by the observation that the transcriptional upregulation of a subset of H2B.8-marked genes during imbibition is affected in absence of H2B.8.

### H2B.8 is dispensable for seed germination

Despite the evolutionary conservation of H2B.8 orthologs in angiosperms, which exhibit conserved expression patterns in seeds and pollen ^26,55^, *h2b.8* mutants do not show altered seed germination or dormancy under our laboratory conditions. Furthermore, altered phenotypes or gene expression defects are not observed upon its ectopic expression under a strong viral promoter ^25,33^ or under the *H3.3* promoter (*our study*). This finding indicates that chromatin plasticity may be sufficient to accommodate H2B.8 loss in seeds or the ectopic presence of this atypical variant along with the potential nucleosome stability changes induced by its incorporation into the chromatin. The absence of a germination phenotype is consistent with the *h2b.8* mutants being fertile, and the specific phenotypic differences observed for the loss of other histone variants under standard growth conditions such as the sperm-specific H3.10^24^ or H3.14^56^ and H3.15 ^57^, which are induced in roots upon stress or wounding. Even though these variants show particular expression patterns, play a role in H3K27me3 reprogramming or stress responses, they are dispensable for plant viability and fertility. This raises the question what drives the conservation of the H2B.8 variant and its specific expression patterns in seeds and pollen through angiosperm evolution. Selection pressure may be primarily exerted on the sperm cells, where H2B.8 incorporation has been shown to reduce the nuclear volume to facilitate migration of the sperm nuclei through the pollen tube ^25^. Alternatively, deficient germination or reduced seed longevity might only be observed in nature, where in contrast to laboratory conditions, seeds are constantly exposed to varying environmental conditions and repeated imbibition-desiccation cycles. Under these conditions, H2B.8 may facilitate the adjustment of the transcriptional program during the process of imbibition. Indeed, when seeds are imbibed under suboptimal conditions and enter secondary dormancy, *H2B.8* expression is induced ^58^ suggesting that the accumulation of H2B.8 can be fine-tuned in response to environmental conditions.

### Eviction of H2B.8 reveals global histone dynamics during imbibition

Upon water uptake, *H2B.8* transcript levels decline rapidly and H2B.8 histone variants are evicted and degraded, a process that involves the proteasome pathway. The degradation occurs first in the cotyledon tissue followed by the hypocotyl, while H2B.8 is retained longer in the root meristem area. This eviction takes place initially in a replication-independent manner and is not restricted to the H2B.8 histone variant. Tagged H3.3 deposited during seed maturation is similarly evicted, revealing global histone turnover in non-replicating cells during the imbibition step. After 48h of imbibition, approximately 50% of the H2B.8 and H3.3 variants that were initially present in dry seeds are evicted. This turnover occurs with similar rates as in alfalfa cell cultures ^59^, yet it contrasts with the substantially longer histone lifetimes of over 100 days seen in mammalian tissue with non-proliferating liver and brain cells ^39^ or to the short histone lifetimes of ∼4 hours in mouse embryonic fibroblasts ^60^. These estimations are global at the cellular or tissue level, but differences among genomic regions depending on their transcriptional activity are expected. Indeed, at genes with high nucleosome turnover, H3.3 half-life ranged from 1-1.5 hours in Drosophila S2 culture cells ^61^ or from 12-24h at transcribed genes in Arabidopsis ^62^. The histone turnover monitored in this study depends on the physiological state of the seed, as evidenced by the accelerated eviction of H2B.8, when seeds are imbibed under continuous light. Subsequent studies should elucidate whether histone turnover during seed imbibition exhibits variation among genomic loci and whether it is driven by transcriptional activity. Global histone turnover is not restricted to the seed-to-seedling transition but has also been observed in the G2 cells of roots during the final cell cycle before the exit from proliferation, where H3.1 is evicted and replaced by H3.3 ^63^ or after fertilization when paternal H3.3 variants are actively removed^29^. Different mechanisms could lead to the observed histone turnover, and our data show a requirement for translation and transcription, suggesting that it is an active process. Global transcriptional restart upon imbibition may favor H2B.8 eviction through the action of *de novo* expressed proteins or enhanced chromatin remodeling activity. A potential outcome of histone turnover is the setting of new post-translational modifications. The replacement of H2B.8, which lacks the C-terminally lysine, by shorter H2B variants that can be ubiquitinylated, may may favor H2Bub enrichment, which has been linked to gene expression reprogramming ^64^.

In this study, we demonstrated that dry seeds are characterized by the enrichment of a specific H2B variant at transcriptionally repressed genes and transposable elements on chromosome arms. The H2B.8 histone variant forms heterotypic nucleosomes and, due to its structural differences, is expected to modify nucleosome structure. Upon imbibition, the H2B.8 and H3.3 histone variants present in the dry seed are evicted, revealing active chromatin reprogramming that initially occurs in a replication-independent manner accompanying transcriptional reprogramming. Whether histone eviction occurs globally or with distinct dynamics at transposons or genes with specific transcriptional dynamics during the seed-to-seedling transition is an exciting avenue for future studies.

## Materials and Methods

### Plant material

The T-DNA insertion mutants *h2b.8-3* (SM_3_38239), *hira-1* (WiscDsLox362H05) and *ssrp1-2* (SALK_001283C) were obtained from the Nottingham Arabidopsis stock center. Plants were grown under long-day conditions (16h light, 8h dark) at 23°C in Aralab growth chambers. Plants were transformed using floral dipping, and transgenic plants selected on Hygromycin, Kanamycin or Basta according to the resistance marker. After-ripened Arabidopsis seeds were used in all experiments, except for dormancy tests. For the imbibition (48h in the dark at 6°C) and germination time-course (long-day conditions, at 23°C), seeds were spread on water-imbibed filter paper in glass petri dishes. Samples were taken every 24h (24hD (dark), 48hD, 48hD24hL(light), 48hD48hL and 48hd72hL). Alternatively, seeds were placed on imbibed filter paper in glass petri dishes directly under continuous light. For inhibitor experiments, a hole was introduced into each individual seed with a fine needle (29G) and seeds incubated in presence of 500μM α-amanitin, 100μM cycloheximidine, 100μM MG132 or a combination of 50μM MG132 and 50μM Bortezomib in a small plastic tube under continuous light. For germination tests, seeds were placed on imbibed filter paper in glass dishes. A seed was scored as germinated as soon as the radicle protruded from the seed coat. Ethylene treatment was carried out by placing seeds into 360 ml gas-tight Petri dishes, in which 0.8 ml gaseous ethylene (5%, Air Liquide, Paris, France) was injected with a syringe to obtain a final concentration of 100 ppm of ethylene.

### Cloning and construction of transgenic lines

Gibson cloning was used to assemble the different constructs. Each fragment was amplified by PCR and inserted into the pBluescript SK-derived vector pTP1 that contains the *att*L1 and *att*L2 Gateway recombination sites (kind gift from T. Pélissier). These constructs were then transferred into Kanamycin-resistance conferring destination vector pGWB401 (Addgene). All constructs were verified by Sanger or whole-vector Nanopore sequencing. To express H2B.8 or H3.3 histones under the control of the *H2B.8* promoter, 1068 bp of the *H2B.8* promoter region and the 5’UTR as well at the 3’UTR region of the *H2B.8* gene were used. The *H2B.8* coding region was cloned in frame with a 4x Myc-tag (*pH2B.8::Myc-H2B.8*) and the *HTR5* coding region with a 3x Flag-tag (*pH2B.8::Flag-H3.3*). To express Myc-H2B.8 under the *HTR5* promoter, 1016 bp of the *HTR5* promoter region, 5’UTR and its 3’UTR were used. For the *pH2B.8::H2B.8-Flag* construct the *H2B.8* promoter, 5’UTR, CDS and 3’UTR region were used and a 3x Flag tag introduced in the C-terminus using the Gibson assembly method. To visualize *H2B.8* and *H2B.4* promoter activity during germination, the *GUS* coding regions were cloned under control of 1068 and 2430 bp long fragments of the *H2B.8* or the *H2B.4* promoters respectively in pDonR221 and then transferred in *pBGWFS7* with Gateway cloning kit. To construct the *pH2B.8::H2B.8-GFP* expressing transgenic plants, 2034 bp of the *H2B.8* promoter region and the 5’UTR as well as the coding region of the *H2B.8* gene were inserted into pENTR1A (Thermofisher). The construct was then transferred into the Basta-resistance conferring destination vector pGWB604. Primers used for cloning are listed in *Supplementary Data 2*.

### X-Gluc histochemical staining

Dissected embryos were fixed for 2h in 80% acetone at-20°C, vacuum infiltrated twice 5 min with 0,5 ml of X-Gluc staining solution (50mM NaPO_4_ pH 7, 20mM EDTA, 0,2% Triton-X-100, 2mM Potassium-Ferrocyanide, 2mM Potassium-Ferricyanure, 2mM XGluc) and incubated 24h at 37°C in the dark. Samples were then repeatedly washed in Ethanol at room temperature (RT) and images acquired with an Olympus Stereo SZH microscope under 15x magnification.

### EdU staining

Approximately 30 seeds were imbibed under agitation in water supplemented with 10µM 5-ethynyl-2’-deoxyuridine (EdU) from the Click-iT® EdU Imaging Kit. The seeds were dissected, and the embryos/seedlings fixed and processed according to the kit’s instructions. Imaging was performed using a spinning disk microscope equipped with a 63x objective.

### Flow cytometry analysis

Fifty seeds were harvested at the different time points, and the embryos were dissected and then chopped in 100µL of cold GB buffer (45mM MgCl₂, 30mM Sodium citrate, 20mM MOPS pH 7, 0.1% Triton X-100, 1% BSA). Samples were then filtered through a 30µm CellTrics filter and fixed in 2% formaldehyde in GB. To purify the nuclei, the filtrate was deposited on top of an Eppendorf tube containing 100µL of 2M sucrose, layered over GSB buffer (0.5M sucrose, 0.1% BSA, 15mM Sodium Citrate, 10mM MOPS pH 7, 0.15% Triton X-100, 0.5% PVP, 0.5% BSA, 22.5mM MgCl₂). Samples were then centrifuged for 4 min at 4000 g with minimal braking. The nuclei, localized in the GSB layer, were collected and diluted in GB buffer with 5µg/mL of DAPI for a final volume of 1mL and analyzed on an Attune NTX Flow Cytometer. Analyses were performed with FlowJo.

### Nuclear image analysis

Dry seed embryos were dissected, stained with Hoechst and imaged as described in^65^. From the 40 central Z-slices of each image, we extracted small 3D images containing one nucleus each, using the ImageJ plugin *NucleusJ* 2.0 ^66,67^. For each genotype, an equal number of nuclei (10-20) per embryo were randomly selected. To segment the nucleus and the intensely stained regions within the nuclei, we generated a deep-learning model using the open-source deep-learning framework *Biom3d* ^67^. In short, *Biom3d* was used to train two separate models, one to segment nuclear volumes, and the other to segment the Hoechst-bright regions and to segment nuclei and intensely stained regions in WT and mutant nuclei. All segmented signals were verified by eye in a double-blind mode after overlay with the respective raw image and obviously aberrant predictions discarded. Quantitative parameters were subsequently extracted from the resulting segmentations. Screenshots of representative nuclei and the predicted segmented nuclei and chromocenter masks were obtained using the 3D view in Napari.

### Immunofluorescence staining

Nuclei were isolated and immunofluorescence experiments carried out as described ^65^. In brief, embryos were finely chopped with a razor blade in 200μl LB01 buffer (15mM Tris-HCl pH 7.5, 2mM NaEDTA, 0.5mM spermine, 80mM KCl, 20mM NaCl and 0.1% Triton X-100) and filtered through a falcon cell strainer cap (40 µm). After a 1:1 dilution in sorting buffer (100mM Tris-HCl pH 7.5, 50mM KCl, 2mM MgCl_2_, 0.05% Tween-20 and 5% sucrose), 20μl of the nuclei suspension were spread on a polylysine slide. After drying, the nuclear suspension was fixed in 2% formaldehyde in 1X PBS for 5 min and rinsed. Slides were washed in 1X PBS +0,5% Triton X-100 for 15min at RT, then 3 times with 1X PBS 5 min at RT. For immunodetection, slides were incubated overnight at 4°C with 20µl of primary antibody in fresh blocking buffer (3% BSA, 0.05% Tween 20 in 1X PBS), washed 3 times for 5 min each in 1X PBS solution and then incubated for 2–3h at RT in 20µl of blocking buffer containing secondary antibodies. Finally, slides were washed 3 times for 5 min each in 1X PBS and mounted in Vectashield mounting medium containing 1.5μg/mL DAPI (Vector Laboratories). The following antibodies were used: anti-Flag (Abcam ab18230, GR229415-5, 1/100), anti-Myc (Millipore 05-724, 3800004, 1/100), anti-H3K9me2 (Abcam ab1220, GR166768-3, 1/100), anti-H2A.Z (Agrisera AS10 718, 2011, 1/250) and anti-H3K27me3 (Diagenode, C154110069, A1818P, 1/100). The anti-H2A.W antibody was generated against two peptides specific for the H2A.W.6 variant as in (Yelagandula et al., 2014) and used to a final dilution of 1/400 in blocking buffer. Primary antibodies were detected with anti-mouse Alexa-594 (1/1000) and anti-rabbit Alexa-488 (1/1000). Images were acquired with an inverted confocal laser-scanning microscope (LSM800; Carl Zeiss). The 488-nm line of a 40-mWAr/Kr laser, and the 544-nm line of a 1-mW He/Ne laser were used to excite GFP/YFP, and RFP, respectively. Images were acquired with a 63x oil immersion objective and stored in an *in-house* server (OMERO) and figures were produced using OMERO figure.

### RNA extraction and RT-qPCR

Total RNA was extracted from dry seeds using a combination of Phenol-Chloroform extraction and RNAzol@RT (Molecular Research Center). In brief, 100mg of seeds were ground in liquid nitrogen using mortar and pestle and taken up in extraction buffer (100mM Tris-HCl pH 9,5; 150mM NaCl, 5mM DTT; 1% Sarkozyl). After vortexing and centrifugation, the supernatant was recovered and mixed with 0.5 Volumes of Chloroform and Acidic Phenol. RNA in the supernatant was precipitated and the pellet taken up in 800mL RNAzol@RT and treated following the manufacturers’ instructions. DNA was removed with DNaseI (Promega) and finally RNA purified using Zymo RNA purification columns (Zymo Research). For *spike-in* RNA seq, 200 embryos imbibed for 24h were manually dissected, harvested into ice cold acetone and ground to fine powder using a metal bead in a Qiagen Tissue Lyser. RNA was extracted using the Qiagen micro kit following the manufacturers’ instructions. After adding the first buffer (RLT PLUS), 100ng of Drosophila total RNA was added to each sample. RNA quality was checked using the Tapestation (Agilent) and quantified by Qubit. Total RNA was reversed transcribed using polyT primers using AMV Reverse Transcriptase (Promega) using the manufacturers’ instructions. Histone *H2B* transcript levels were normalized to *MON1 (AT2G28390)* and *TIP41 (AT4G34270)* using the Roche Lightcycler and qPCR kit (*Supplementary Data 2*).

### RNA-seq analysis

Library preparation from total RNA and mRNA sequencing was performed by BGI. For dry seeds, raw reads were processed by the bioinformatics pipeline CRESCENT (https://github.com/gilless429/crescent) developed with the snakemake workflow management system. Quality checking was done using fastqc and multiqc. Low-quality reads were trimmed or removed by cutadapt. Reads were then mapped to the TAIR10 reference genome via the hisat2 mapper, before being assigned to genomic features by featureCounts.

Differential expression analysis was executed through the DESeq2 R package and read counts were normalized using DESeq2’s median of ratios method. The thresholds for what was considered a mis-regulated feature were an adjusted p-value (Benjamini-Hochberg procedure) of less than 0.05 and a log2FC of absolute value greater than 1. The PCA was produced using DESeq2’s plotPCA, heatmaps using the pheatmap package, after variance stabilizing transformation of read counts and Venn diagrams with the ggvenn package.

For *spike-in* RNA seq analysis of 24h imbibed seeds, the same snakemake pipeline was used. Reads were mapped to an “hybrid” genome containing the *Arabidopsis thaliana* TAIR10 reference genome and the *Drosophila melanogaster* reference genome DM6 by hisat2 ^68^ mapper, processed and deduplicated with samtools. Mapped reads were then separated depending on their origin. Arabidopsis reads were assigned to Arabidopsis genomic features by featureCounts. We calculated for each sample an α factor corresponding to the percentage of Drosophila reads divided by the number of Arabidopsis reads. The *spike-in* factor for each sample were calculated by dividing the α factor of each sample by the smallest α factor. Differential expression analysis was executed through the DESeq2 R package within CRESCENT and read counts were normalized using DESeq2’s median of ratios method.

### Chromatin-Immunoprecipitation followed by sequencing

100mg of seeds expressing the Myc-H2B.8 protein under control of its endogenous promoter were ground to a fine powder in liquid nitrogen cooled mortars. The powder was taken up in Extraction buffer (EB1, 0.4M sucrose, 10mM Tris-HCl, pH8, 10mM MgCl_2_, 10mM β-mercaptoethanol, 0.1mM PMSF, proteinase inhibitors (Roche)) and the solution filtered through two layers of miracloth. After centrifugation (20 min at 3 000 g at 4°C), the pellet was taken up in EB2a (0.25M sucrose, 60mM Hepes, pH8, 10mM MgCl_2_, 0.3% Triton X-100, 0.1mM PMSF, proteinase inhibitors (Roche)) and Formaldehyde added to a final concentration of 1%. After 10 min incubation, crosslinking was stopped by adding glycine (140 mM final). Formaldehyde was removed by one wash in EB2a and one in EB2B (0.25M sucrose, 10mM Tris-HCl, pH8, 10mM MgCl_2_, 1% Triton X-100, 5mM β-mercaptoethanol, 0.1mM PMSF, proteinase inhibitors (Roche)). The nuclear pellet was taken up in 100ul of Nuclei Lysis (NL) buffer (50mM Tris-HCl, pH8, 10mM EDTA, 1% SDS, 0.1mM PMSF, and proteinase inhibitors (Roche)) and left in the fridge for 1.5h. After dilution of with NL buffer without SDS to 0.1% SDS final, chromatin was shared using a Covaris sonicator (IP 140, CBP 200, DF 10 %, T° 5°C, 6min). Chromatin was stored overnight in the fridge to allow checking sonication efficiency by decrosslinking of a chromatin aliquot overnight and DNA purification using ChIP DNA Clean & Concentrator columns (Zymo Research). Chromatin was incubated with 2µl of either mouse monoclonal anti-Myc (Millipore, 05-724) or rabbit polyclonal anti-Myc (Abcam, AB9106) overnight, before adding 30µl of Dynabeads and incubating for at least 3 hours. Beads with chromatin were washed in low-salt and high-salt buffers as previously described ^69^ before elution in 0.1M NaHCO_3_ and 1%SDS. Chromatin was decrosslinked overnight, treated with Proteinase K and RNase A before purification using ChIP DNA Clean & Concentrator columns (Zymo Research). Several IPs were pooled for either antibody using a ChIP DNA Clean & Concentrator column and DNA concentration determined using Qubit. Library preparation and sequencing was carried out by BGI.

### ChIP-seq analysis

After reads quality check with Fastqc, reads were aligned to the TAIR10 genome using Bowtie2^70^. PCR duplicates were discarded using samtools rmdup ^71^. Peaks were called using MACS2 (--qvalue 0.01,--bw 300) using corresponding inputs as background, only peaks overlapping by at least 10% between replicates were retained. Sample tracks were generated using deeptools bamCoverage ^72^ with option-normalizeUsing RPGC. Based on H2B.8 peaks’ fold change compared to Input calculated by MACS2 ^73^, H2B.8 peaks were divided in 3 quantiles with high, middle, and low H2B.8 levels to investigate the impact of H2B.8 on chromatin interactions. To compare H3K4me3 and H3K27me3 enrichment at the H2B.8 gene in seeds and seedlings (data from ^30,74^) data were normalized using S3norm ^75^. All datasets used in this study are listed in *Supplementary Data 3*.

### Hi-C

In situ Hi-C experiments were performed using 400 manually dissected WT and *h2b8-3* mutant dry seed embryos per replicate according to ^76^ using the DpnII restriction enzyme (NEB). Likewise, the second crosslinking step was performed involving disuccinimidyl glutarate and a second digestion with DdeI (NEB) according to ^77^. DNA libraries were prepared using NEBNext Ultra II DNA library preparation kit (NEB) according to the manufacturer’s instructions (10 cycles for the PCR amplification step). DNA libraries were checked for quality and quantified using a 2100 Bioanalyzer (Agilent) and the libraries were subjected to 2 × 75 bp high-throughput sequencing by NextSeq 500 (Illumina). Two independent biological replicates were generated yielding over 39 million valid interaction pairs for both control and mutant. The proportion of cis (intrachromosomal) interactions, an indicator of Hi-C data quality, was 85% for the control and 83% for the mutant condition, consistent with the values reported for *in situ* Hi-C in Arabidopsis confirming the robustness and reliability of our Hi-C datasets.

### Hi-C analysis

Raw Illumina sequencing fastq files were processed with Trimmomatic v0.38 ^78^. Cleaned reads were mapped independently to the Tair10 reference genome (available at https://www.ncbi.nlm.nih.gov/datasets/genome/GCF_000001735.3/) using HiC-Pro v3.0.0 ^79^.

Bowtie2 parameter settings were default, except “BOWTIE2_GLOBAL_OPTIONS =--very-sensitive-L 30--score-min L,-0.6,-0.6--end-to-end –reorder” and “BOWTIE2_LOCAL_OPTIONS =--very-sensitive-L 20--score-min L,-0.6,-0.6--end-to-end--reorder” Forward and reverse mapped reads were paired and assigned to DpnII and DdeI restriction fragments. Invalid ligation reads, such as dangling ends, were filtered. Then HiC-Pro v3.0.0 with parameter “-s merge_persample” was performed to merge each pair of replicates before using the valid pairs to generate interaction. After, ‘.hic’ files were generated with the hicpro2juicebox.sh script, which belongs to HiC-Pro then furtherly transformed to ‘.cool’ files and ‘.h5’ files using the function hicConvertFormat from HiCExplorer ^80^ at different resolutions. Then, the’.h5’ files were normalized using HiCExplorer functions, specifically’hicNormalize’ and’hicCorrectMatrix’ with the’--correctionMethod KR’ parameter. The scaling plot of Interaction frequencies against genomic distance was drawn at a 100 kb resolution from the’.h5’ files using’hicPlotDistVsCounts.R’ ^81^. The saddle plots of chromatin compartmentalization and compartment strengths boxplot were drawn at 20kb resolution by‘CompareTwoHiCmatrixV2BasedOnHiCdat.R‘ ^81^. The figures of aggregated Hi-C sub-matrices of different H2B.8 peak quantiles are plotted by coolpup.py ^82^ at 25kb resolution.

### Nuclear protein extraction and Western Blot

70 milligrams of dry or imbibed seeds were frozen and ground in liquid nitrogen to a fine powder and homogenized in Honda’s buffer. Nuclei isolation was carried out as previously described ^83^ with minor modifications. Nuclear pellets were resuspended in 100μl of Laemmli 4X buffer, supplemented with β-mercaptoethanol (Bio-Rad), incubated for 5 min at 90°C and centrifuged at 16000 g for 5 min. 15-20μl of nuclear protein extracts were loaded on 4-20% precast SDS-PAGE gels (Bio-Rad) for Western Blot analysis, using standard procedures. Western Blots were probed with different primary antibodies: monoclonal anti-Myc antibody (Millipore, 05-724, 1/2000), polyclonal anti-Myc antibody (Abcam, AB9106, 1/1000), anti-human H2B antibody (Abcam, ab1790, 1/2000), and anti-H3 antibody (Abcam, AB1791, 1/10 000). The anti-H2B.8 antibody was raised in rabbits (Eurogentec) against an AtH2B.8 N-terminal peptide (VSVTKKKKVVEETIK). The antiserum was further purified by peptide affinity column. Primary antibodies were revealed by incubation with a horseradish peroxidase-coupled anti-rabbit or anti-mouse secondary antibody (Abliance, 1/5000). Immunoblot chemiluminescence was revealed using ECL protein gel blotting detection reagents (Clarity Western ECL Substrate, Bio-Rad, France). Densitometric analysis of immunoreactive protein bands was performed on non-saturated signals, using Image Lab software (Bio-Rad, France).

### Immunoprecipitation of mononucleosomes

200 milligrams of dry seeds (from WT and from Myc-tagged H2B.8 lines) were ground in liquid nitrogen to a fine powder. The powder was solubilized in 15ml of extraction buffer (10mM Tris-HCl pH8, 0,25M sucrose, 10mM MgCl_2_, 1% Triton X-100, 5mM β-mercaptoethanol, supplemented with proteases inhibitors from Roche), followed by vortexing until a fine suspension was obtained. After incubation for 15 min on ice, the homogenate was filtered through one layer of Miracloth (Merck, Millipore) and nuclei were pelleted by centrifugation at 1400g for 15 min at 4°C. Nuclei were resuspended in 10mL of extraction buffer by pipetting up and down. The nuclei suspension was filtered once again, through one layer of Miracloth and the filtered solution centrifuged at 1400 g for 15 min at 4°C. The nuclear pellet was washed once in 1mL of MNase digestion buffer (10mM Tris-HCl, pH7,5, 15mM NaCl, 60mM KCl, 1mM CaCl_2_ supplemented with protease inhibitors). After centrifugation at 1400g for 15 min at 4°C, isolated nuclei were resuspended in 250μl of MNase digestion buffer. Approximatively 350μl of suspension was obtained. An aliquot of 50μl was kept on ice, as a control of non-digested chromatin. 12 U of MNase (Takara) were added to the nuclei suspension before incubation for 30 min at 37°C. During the incubation, nuclei were mixed gently every 5 min. At the end of incubation, an aliquot of 50μl was removed to check digestion efficiency. MNase digestion was stopped on ice by addition of 28μl of STOP solution (100mM EDTA, 100mM EGTA) and nuclei were lysed by addition of 30μl of NaCl 5M (final concentration of 500mM). The suspension was mixed by inverting the tubes and then kept on ice for 15 min. Extracts were cleared by centrifugation for 10 min at 20,000g at 4°C. Supernatant (approximatively 300μl) was diluted by adding 700μl of MNase buffer. An aliquot of 50μl was removed for the input. 5μl of monoclonal anti-Myc antibody (Millipore, 05-724) diluted in 200μl of PBS 1X 0,1 % Tween, was bound on 50μl of protein G magnetic beads (Dynabeads, Invitrogen) by incubating on a wheel for 30 min at RT. Beads were then incubated with MNase extracts (1mL) over night at 4°C on a wheel. Beads were washed 3 times with 200μl of PBS 1X 0,1 % Tween. Then, beads were resuspended in 30μl of Laemmli 4X buffer supplemented with β-mercaptoethanol (Bio-Rad) and they were incubated for 5 min à 90°C and centrifuged at 16000 g for 5 min. 5-10μl of immunoprecipitated proteins were loaded on 4-20% precast SDS-PAGE gels (Bio-Rad) for Western Blot analysis using standard procedures. To check if mononucleosomes were obtained after MNase digestion, DNA was extracted from aliquots of both nondigested and MNase-digested nuclei. 50μl of STOP solution (100mM EDTA, 100mM EGTA) with 1,5μl of proteinase K (Invitrogen, 20mg/mL) were first added to each aliquot before incubation for 30 min at 37°C. DNA was then extracted with CTAB buffer (2% CTAB, 100mM Tris-HCl pH8, 1.5M NaCl, 20mM EDTA pH8) and isoamylic alcohol/chloroform, before precipitation with isopropanol. DNA was taken up in TE buffer (10mM Tris-HCl pH7.5, 5mM EDTA) and treated by RNAse A / T1 (Invitrogen) for 30 min at 37°C. The DNA concentration was determined using a Nanodrop and equal amounts for each sample were run on a 1,5% agarose gel.

## Data availability

RNA-seq, ChIP-seq and Hi-C data are deposited under the record <u>GSE304463</u>. Source data underlying Fig. 3b, c and Figure S6, images used to train deep learning segmentation models and the corresponding models are accessible here: https://omero.mesocentre.uca.fr/webclient/?show=project-2803.

## Acknowledgements

We acknowledge support from ANR grants 4D-Heat (ANR-21-CE20-0036, to MB), SeedChrom (ANR-22-CE20-0028, to AVP), EpiLinks (ANR-22-CE20-0001, to SA), and NUCLINV (ANR-24-CE20-5404, to CT) and support from the French government IDEX-ISITE initiative 16-IDEX-0001 (CAP 20-25). SP is supported by the MSCA DN *EpiSeedLink* (Grant number: 101073476). K.T. acknowledges support from MEXT KAKENHI (grant numbers JP23H04205 and 22K06269) and from the Human Frontier Science Program (RGP0009/2018) from the International Human Frontier Science Program Organization. AVP, CT, MB and CB acknowledge networking support from the GDR EPIPLANT. We thank S. Swiezewski for advice with the transcription inhibitor experiments and the Clermont Imagerie Confocale (CLIC, iGReD, Clermont Auvergne University) for help with confocal microscopy and image analysis.

## Author contributions

LS and AVP conceived the study. LS, SP, MV, SC, SLG, DL, AMC, SD and KT performed experiments and analyzed data. QW, GS, SA and LS analyzed Hi-C, RNA-seq and ChIP-seq datasets. AVP, CB, MB, KT and CT acquired funding. AVP wrote the manuscript with support from all authors.

---

**Supplementary Figure 1:**
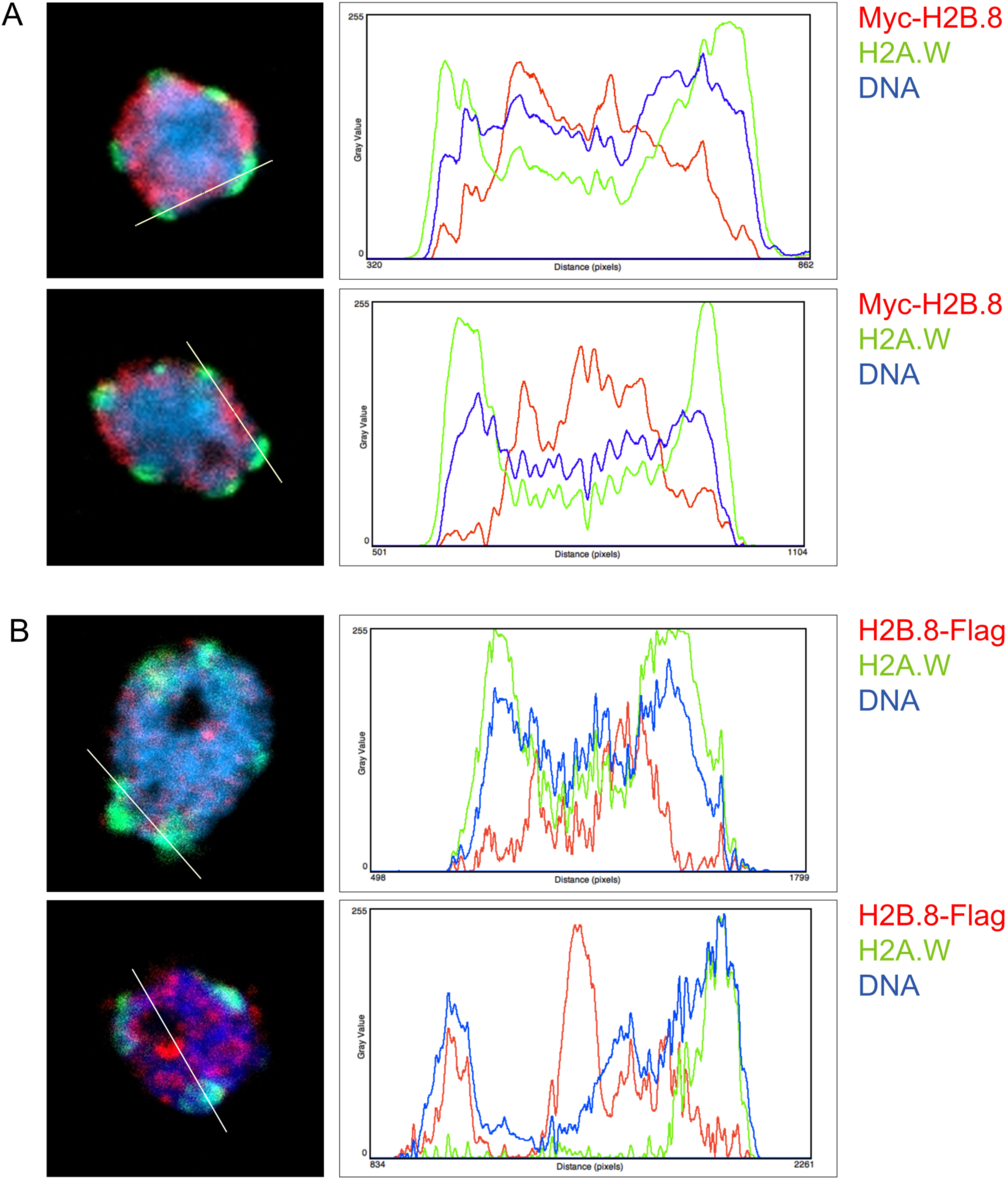
H2B.8 is depleted from H2A.W-marked heterochromatin. Single Z-slices of DS nuclei expressing Myc-H2B.8 (**A**) or H2B.8-Flag (**B**) stained with anti-H2A.W (green), anti-Myc (red) or anti-Flag (red) antibody and counterstained with DAPI (blue). ImageJ was used to draw lines and plot intensity profiles. Most of the chromatin regions that stain intensely with DAPI are marked by H2A.W and therefore correspond to pericentromeric heterochromatin.

**Supplementary Figure 2:**
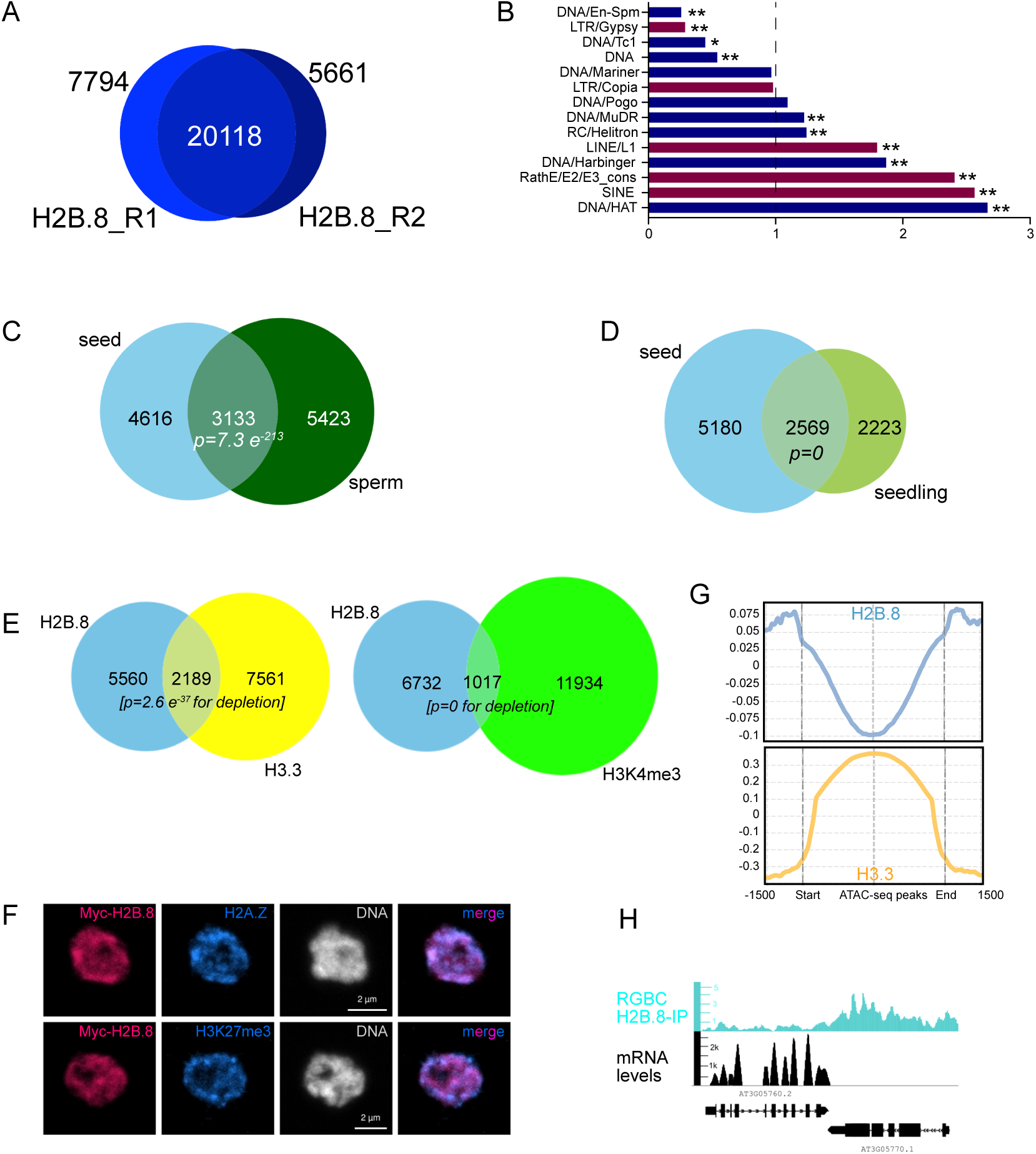
Overlap between H2B.8-marked genes in seeds, sperm and ectopically expressing seedlings and depletion from H2K4me3-marked genes. **A.** Venn-diagram showing overlap between H2B.8 peaks determined by MACS2 in the two Myc-H2B.8 ChIP-seq replicates from dry seeds. Peaks from the two biological replicates, in which Myc-H2B.8 was either immunoprecipitated with a monoclonal or a polyclonal anti-Myc antibody, show substantial overlap. **B.** H2B.8 occupancy at transposable elements (n= 8713) for different transposon classes. DNA transposons are shown in blue and retrotransposons in violet. Odd Ratio > or < 1, \**p <0.05* and *\*\*p<0.0005*, Fishers Exact test. **C** - **D.** Venn diagrams depicting overlap between H2B.8-marked genes in seeds and sperm cells (**C**) or seedlings overexpressing H2B.8 from the 35S promoter (**D**, ChIP-seq data were obtained from ^25^). Significant overlap was assessed with a hypergeometric test. **E.** H2B.8-marked genes are significantly underrepresented among genes enriched in H3.3 (hypergeometric test, data from ^27^) or H3K4me3 (data from ^30^). **F**. Representative DS embryo nuclei stained with DAPI (grey), in which H3K27me3 and H2A.Z (blue) were revealed by immunostaining together with H2B.8 as a N-terminal Myc fusion (magenta). **G.** Metagene plot showing H2B.8 and H3.3 enrichment profiles centered on ATAC-seq peaks in dry seeds (from ^9^). **H.** Genome browser view of an H2B.8-depleted and enriched gene in DS together with the corresponding RNA-seq expression profile.

**Supplementary Figure 3:**
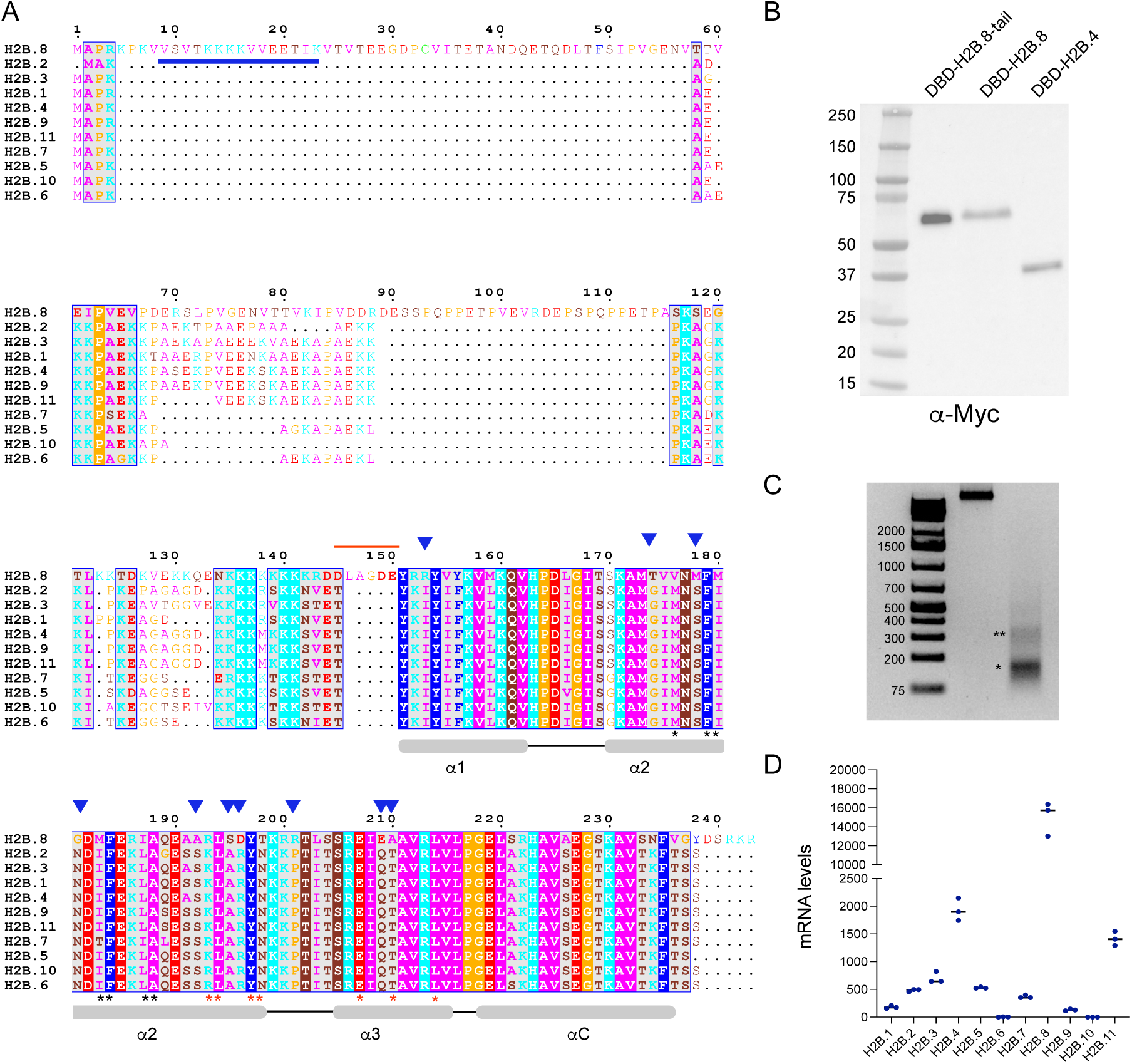
Amino-acid differences between H2B variants and H2B.8 expression in seeds. **A.** Alignment of the 11 Arabidopsis H2B proteins generated with MUSCLE and ESPript 3.0. Residues are colored as follows: HKR (positively charged), cyan; DE (negatively charged), red; STNQ (polar), maroon; AVLIM (hydrophobic), pink; FYW, blue; PG orange; C, green. Positions of the three alpha helices (*α*1, *α*2 and *α*3) constituting the histone fold domain and the C-terminal alpha helix (αC) of all H2B histones (except H2B.8) are indicated in grey. Critical amino acids in the α2 and α3 helices of H2B involved in interactions with H2A α2 or in the H4-H2B four-helix bundle (as described in ^35^) are indicated by black and red stars respectively. Blue arrows indicate differences in amino acids in the α2 and α3 helices between H2B.8 and the other H2B variants including one residue involved in the H4-H2B four-helix bundle (210A). The red bar indicates an H2B.8-specific region, comprising two additional acidic amino acids. The blue bar in the H2B.8 N-terminal tail indicates the epitope used to generate the H2B.8-specific antibody. **B.** Western Blot of total yeast proteins expressing H2B.4, H2B.8 and only the H2B.8 tail (AA1-150) as a fusion with the Gal4 DNA Binding domain (DBD). DBD-histone fusion proteins carrying the Myc tag were detected with an anti-Myc antibody. Molecular weights of 38.3 kDa for Gal4-H2B.4, 49.1 kDa for Gal4-H2B.8, and 38.5 kDa for Gal4-H2B.8-Tail were predicted. **C.** Agarose gel showing undigested DNA (middle lane) and DNA from MNase digested chromatin (right lane) of dry seed nuclei used in nucleosome immunoprecipitation, revealing almost complete digestion to mono-nucleosomes (*), small amounts of dinucleosomes remain visible (**). **D.** Transcript levels (DESeq2 normalization) of the 11 *H2B* genes determined by RNA-seq in *A. thaliana* WT dry seeds.

**Supplementary Figure 4:**
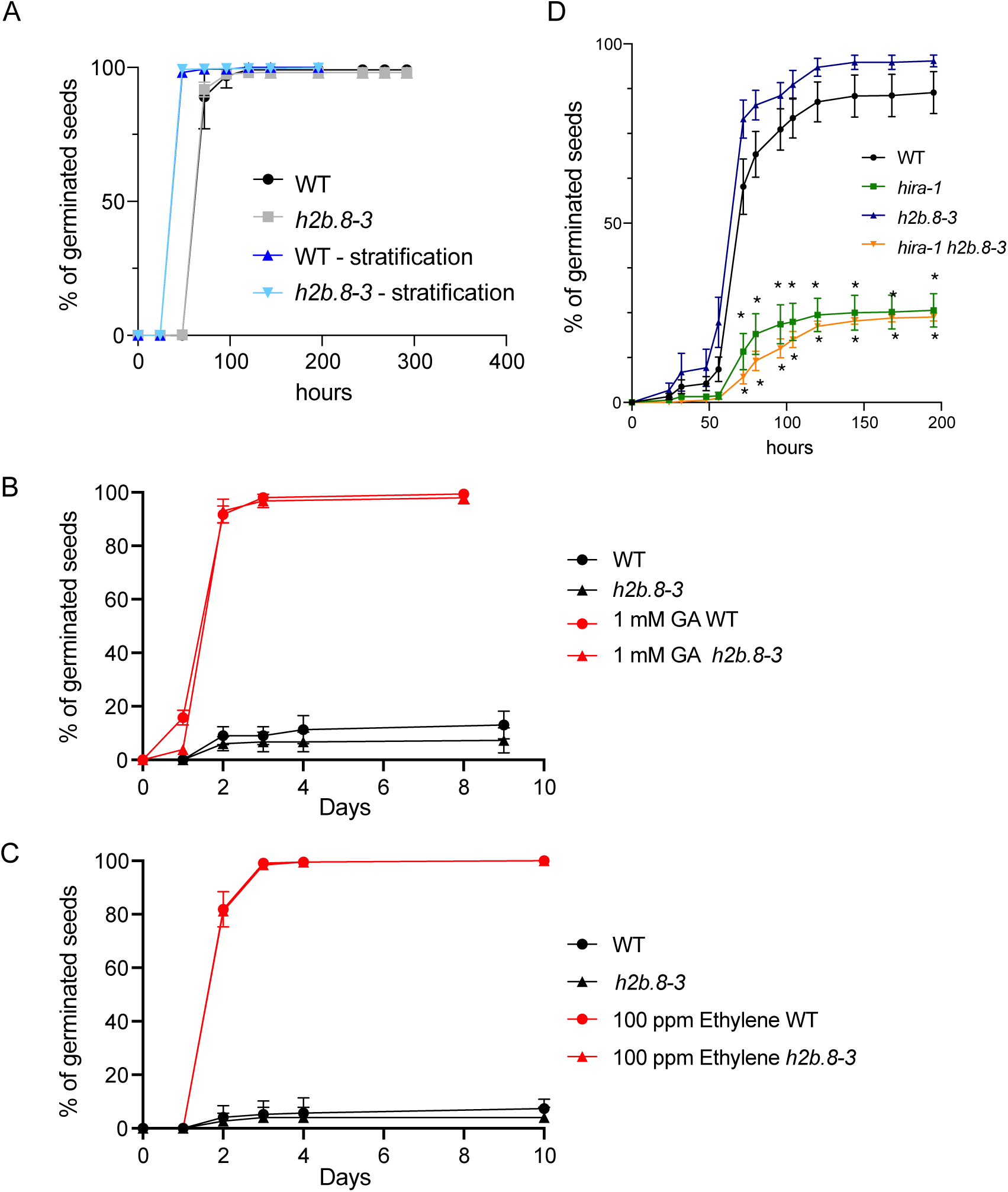
Seed germination is unaffected by the absence of H2B.8. **A.** Seed germination of after-ripened stratified and non-stratified WT and *h2b.8-3* mutant seeds. Mean of 6 biological replicates for each genotype are shown. Error bars correspond to standard deviation. **B-C.** Germination of after-ripened WT and *h2b.8-3* mutant seeds in the absence or presence of gibberellin (GA, 1mM, **B**) and Ethylene (100ppm, **C**). Germination tests were performed at 25°C in the dark. Mean of 3 biological replicates for each genotype are shown. Error bars correspond to standard deviation. **D.** Germination of freshly harvested and non-stratified seeds in WT, *h2b.8-3*, *hira-1* and *hira-1 h2b.8-3* double mutants at 23°C in the dark. Error bars show SEM from 5 or 6 biological replicates. Stars indicate statistically significant differences compared to the WT, *p < 0.005*, 2way ANOVA coupled to a Tukey’s multiple comparisons test.

**Supplementary Figure 5:**
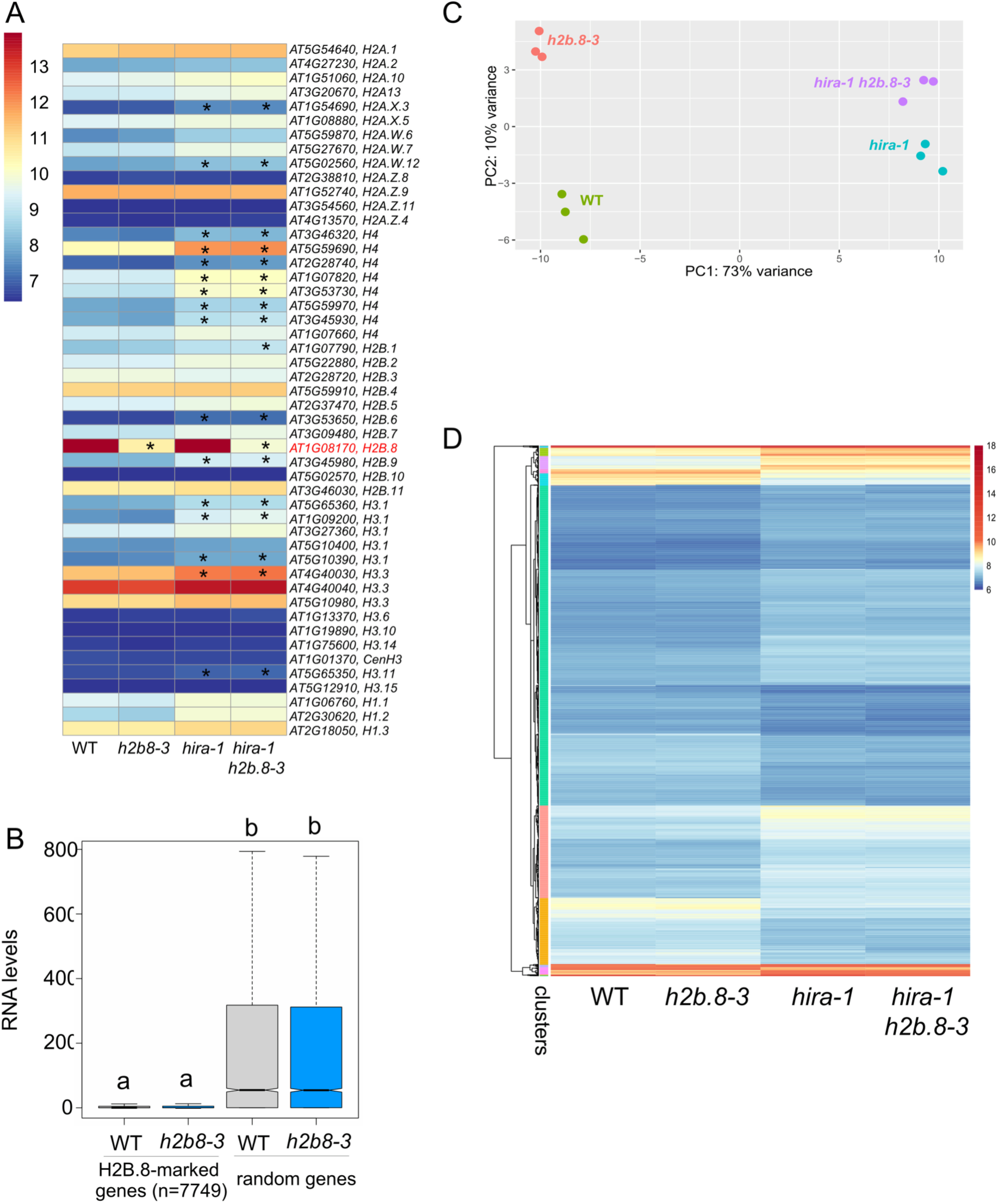
Gene expression changes upon depletion of H2B.8 and/or HIRA in dry seeds. **A**. Heatmap showing normalized read counts of histone genes in WT, *h2b.8-3*, *hira-1* and the *hira-1 h2b8-3* double mutant dry seeds. Stars indicate significantly up or down-regulated genes (log2FC > 1, padj < 0,05) compared to the WT. **B**. Transcript levels (DESeq2 normalization) of an equivalent number (n=7549) of random and all H2B.8-marked genes in WT and *h2b.8-3* mutant dry seeds, significant differences indicated by letters were determined with a Kruskal Wallis test followed by a Dunn’s multiple comparison test, *p < 0.0001*. **C**. Principal component analysis of the three RNA-seq replicates for WT, *h2b.8-3*, *hira-1* and *hira-1 h2b.8-3* double mutants. Single *hira-1* and *hira-1 h2b.8-3* double mutants cluster together. **D**. Heatmap showing normalized read counts for all differentially regulated genes in the dataset.

**Supplementary Figure 6:**
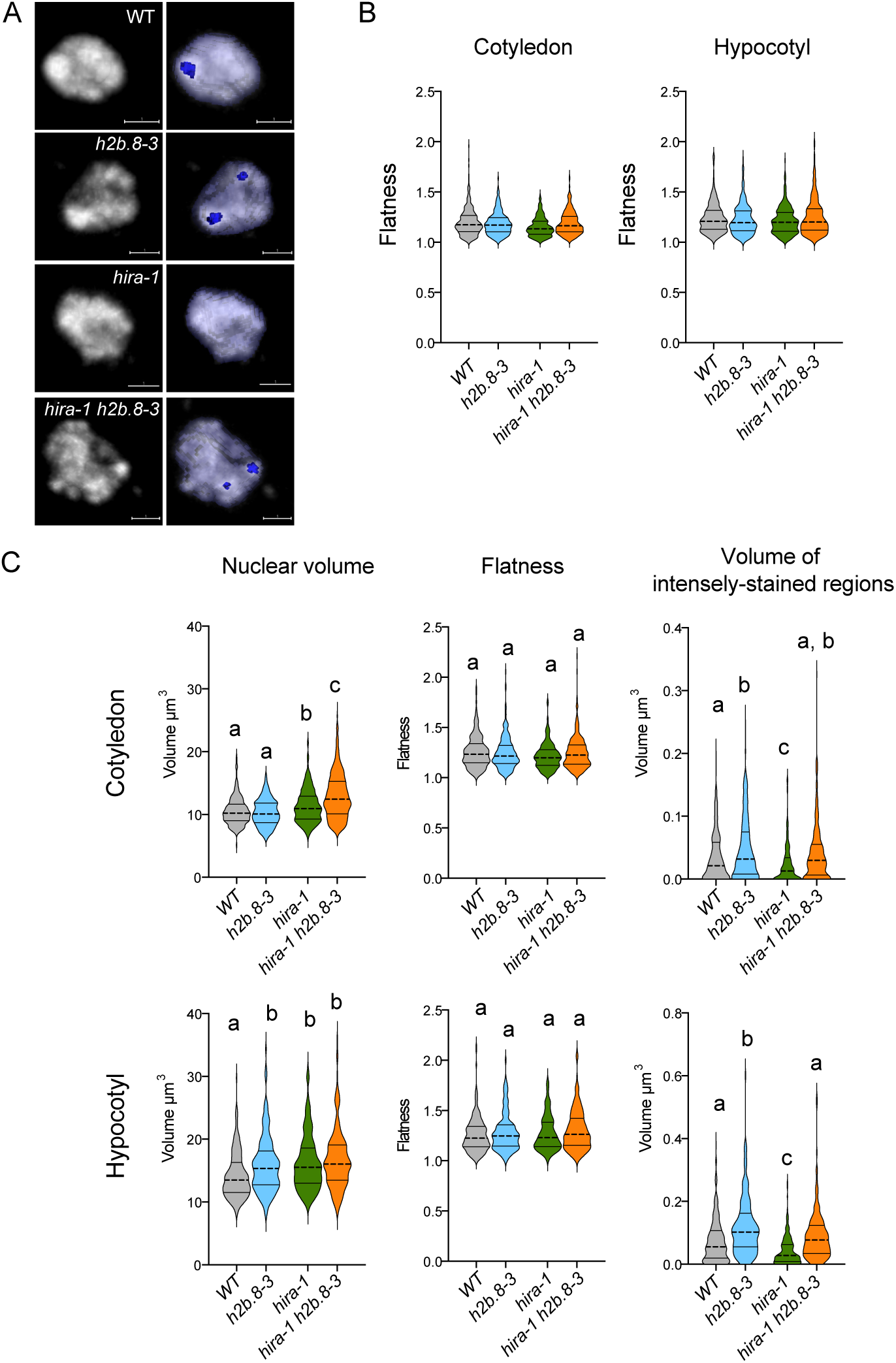
Changes in nuclear and chromatin organization upon depletion of H2B.8 and/or HIRA. **A**. Representative dry seed cotyledon nuclei from WT, *h2b.8-3*, *hira-1* and *hira-1 h2b.8-3* double mutants, in which the nucleus (light blue) and intensely stained chromatin regions (dark blue) were segmented using a deep-learning model developed in *biom3d*. Images were generated with Napari. **B**. Nuclear flatness in cotyledon and hypocotyl nuclei in WT and the three mutant genotypes (*h2b.8-3*, *hira-1* and *hira-1 h2b.8-3*). Nuclei belong to the dataset in Figure 3b and c. **C**. Second independent dataset from an independent seed lot, showing nuclear volume, flatness of the nucleus, and total volume of intensely stained chromatin regions per nucleus. The parameters were determined from cotyledons and hypocotyls from dissected DS using the same deep-learning model generated in *biom3d* as in Figure 3b. Cotyledons: WT: n (number of nuclei) = 233, N (number of embryos) = 24, *h2b.8-3*: n= 273, N =19; *hira-1*: n= 268, N= 19 and *hira-1 h2b.8-3*: n= 271, N= 19. Hypocotyls: WT: n = 229, N = 24, *h2b.8-3*: n= 166, N =19; *hira-1*: n= 174, N= 19 and *hira-1 h2b.8-3*: n= 179, N= 19. Significant differences between were determined using an ordinary one-way ANOVA coupled to a Tukey’s multiple comparisons test.

**Supplemental Figure 7:**
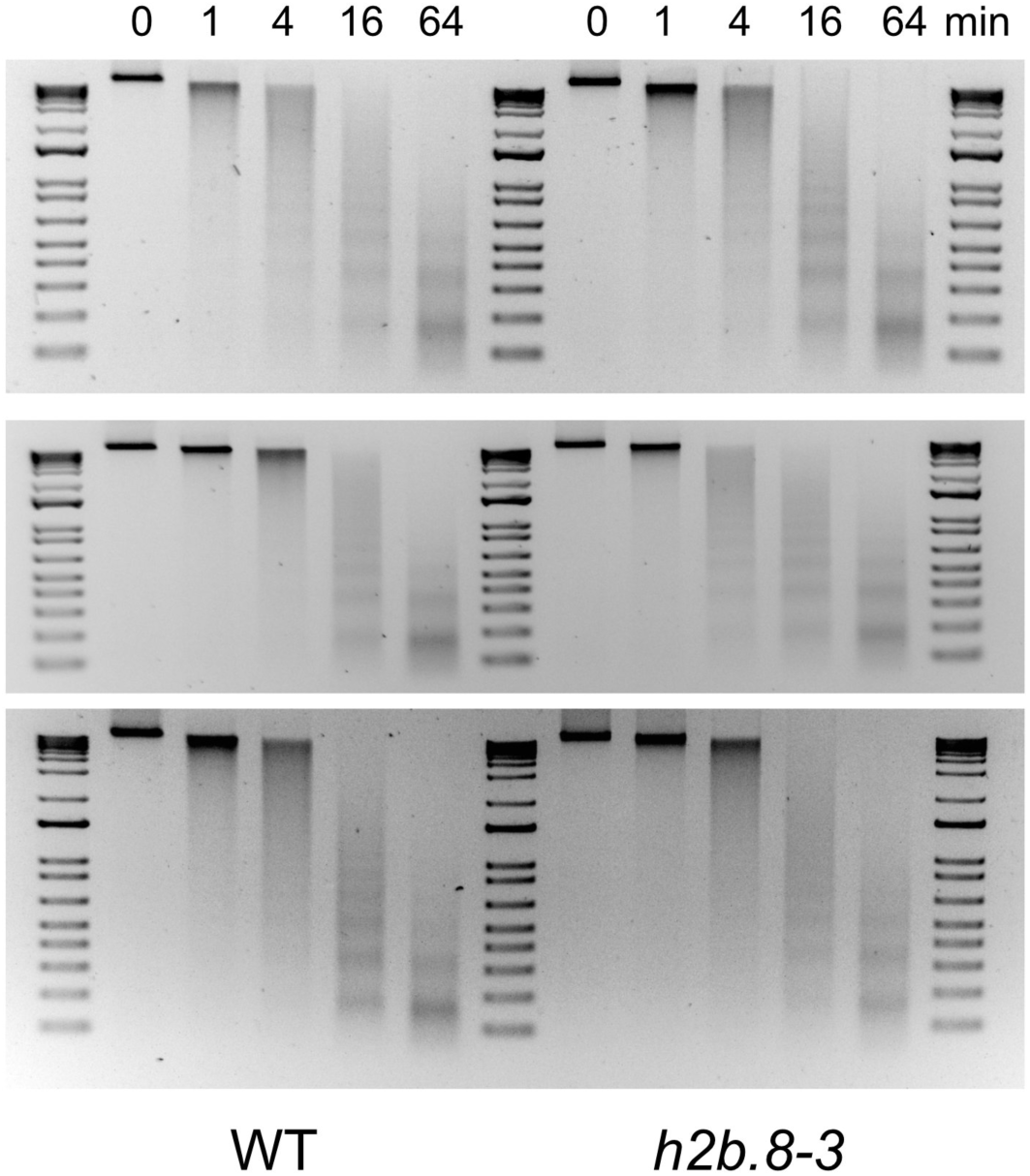
Global chromatin accessibility is unaffected in h2b.8-3 mutant dry seeds. Chromatin accessibility in WT and *h2b.8-3* mutant dry seeds was assessed by MNase digestion for different durations in three independent biological replicates from different seed batches.

**Supplementary Figure 8:**
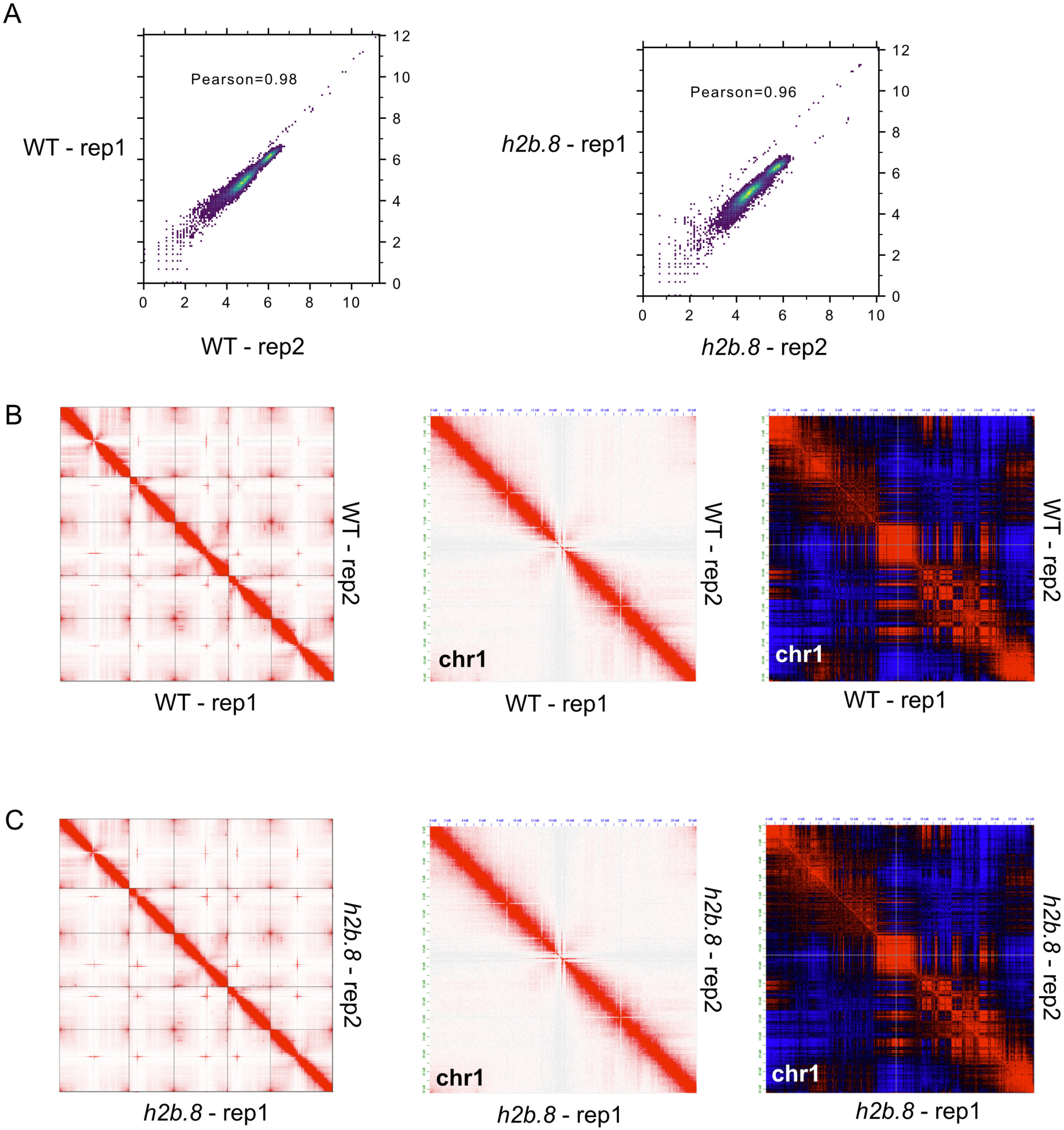
Correlation between Hi-C contact matrices. **A**. Scatter plots showing correlations between Hi-C contact matrices of the two dry seed WT replicates (left) and the two *h2b.8* replicates (right). **B, C**. Hi-C heatmaps of interaction frequencies for the two biological replicates of WT (**B**) and *h2b.8* mutant DS (**C**), in all chromosomes (left), and in chromosome 1 (middle), as well as using the Pearson scaling method in chromosome 1 (right).

**Supplemental Figure 9:**
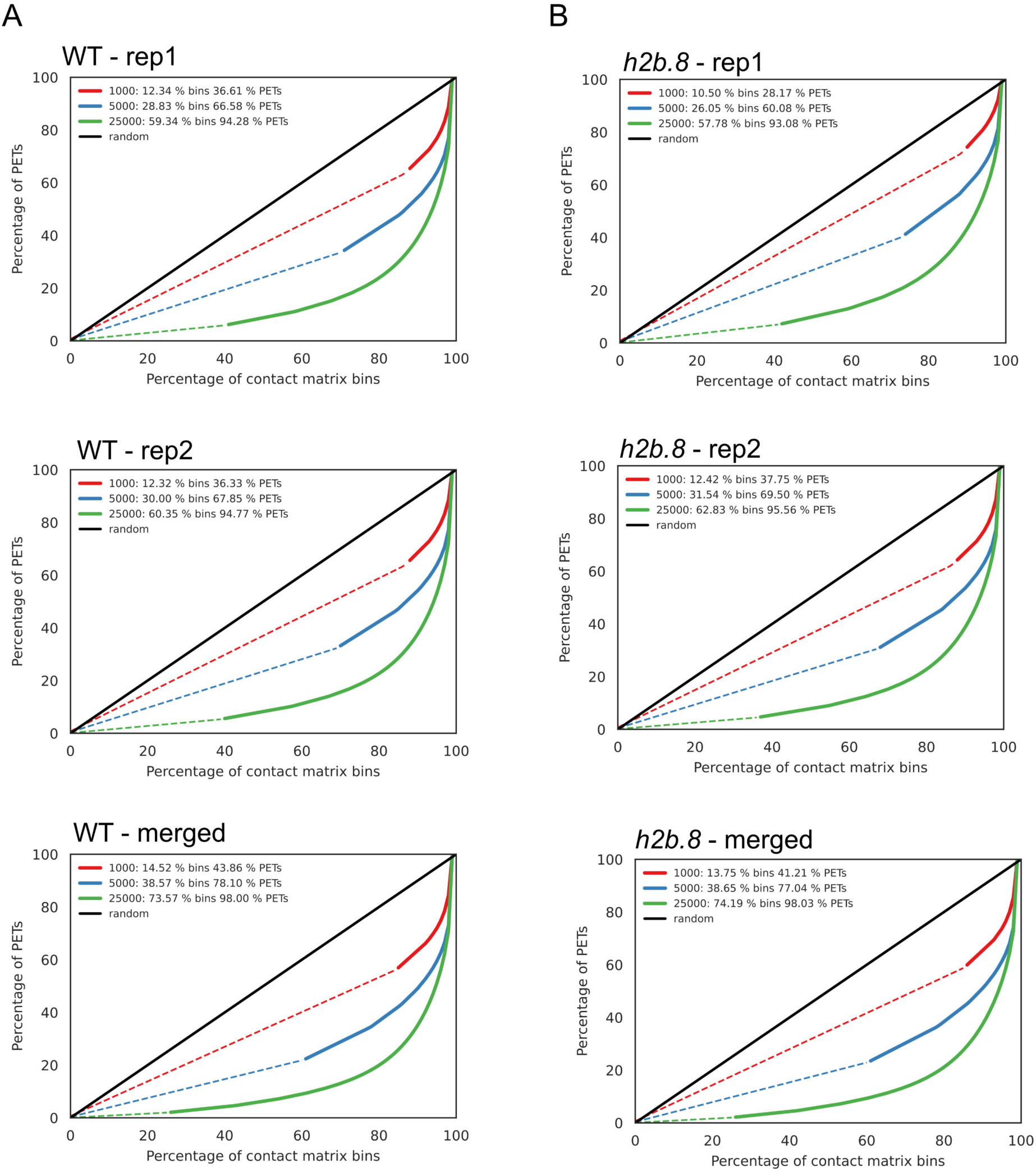
**Estimation of the contact matrix resolution A-B**. Line plots showing the percentage of paired-end tags (PETs) on the percentage of contact matrix bins for WT (**A**), and the *h2b.8* mutant (**B**). The dashed lines indicate the bins from the contact matrix with only singleton PET, and the solid lines indicates the bins with multiple PETs. The random line indicates all PETs are evenly distributed in the contact matrix bins.

**Supplemental Figure 10:**
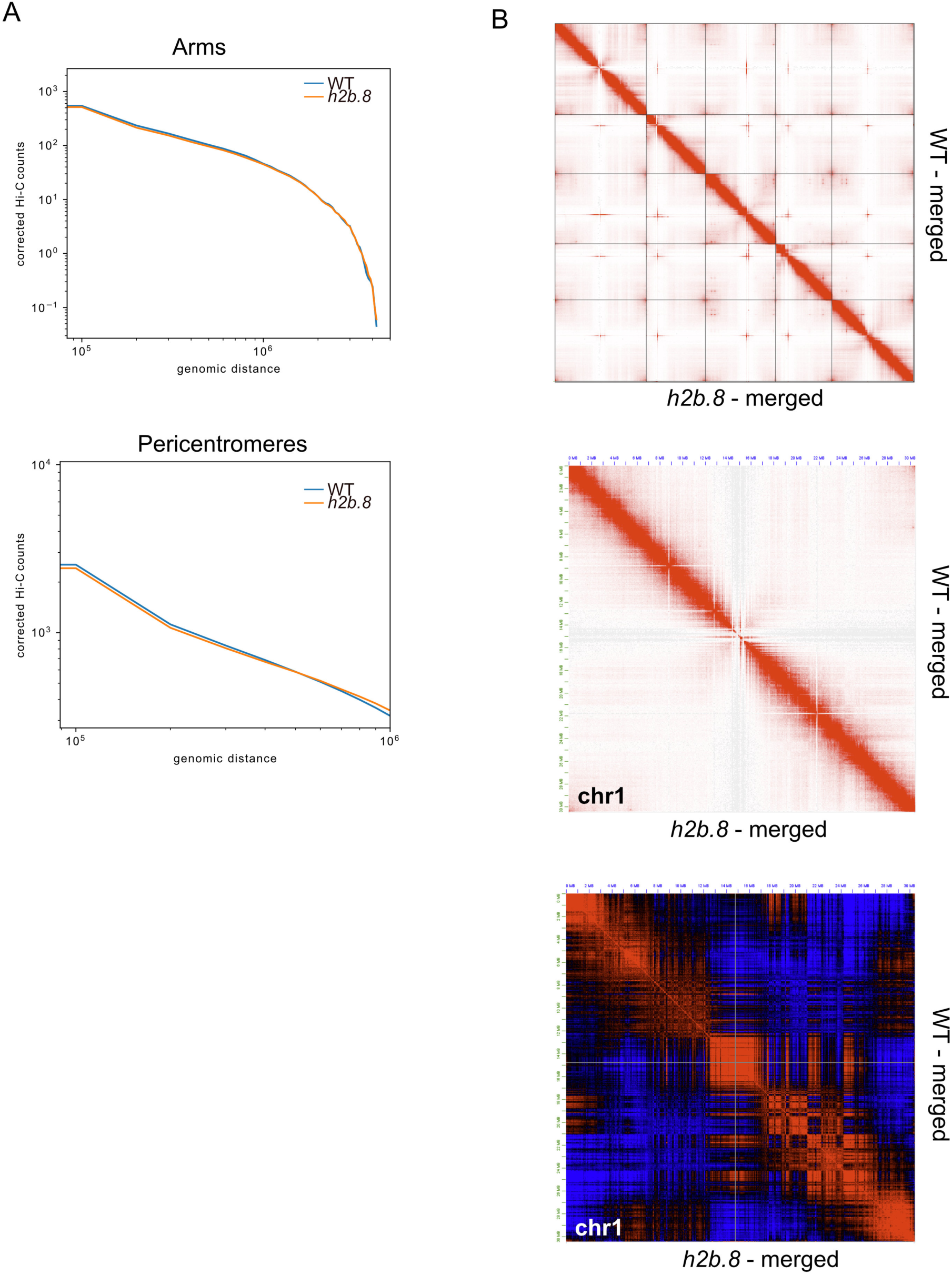
Hi-C heatmaps of interaction frequencies. **A**. Averaged scaling plots with a 100 kb genomic bin size for WT and *h2b.8* mutant dry seed embryos, illustrating how interaction frequencies vary with increasing genomic distance for all chromosomes on chromosome arms (top) and pericentromeric regions (bottom). **B.** Hi-C heatmaps of interaction frequencies for merged WT samples and merged *h2b.8* mutant samples, in all chromosomes (top), and in chromosome 1 (middle), as well as using Pearson scaling method in chromosome 1 (bottom).

**Supplementary Figure 11:**
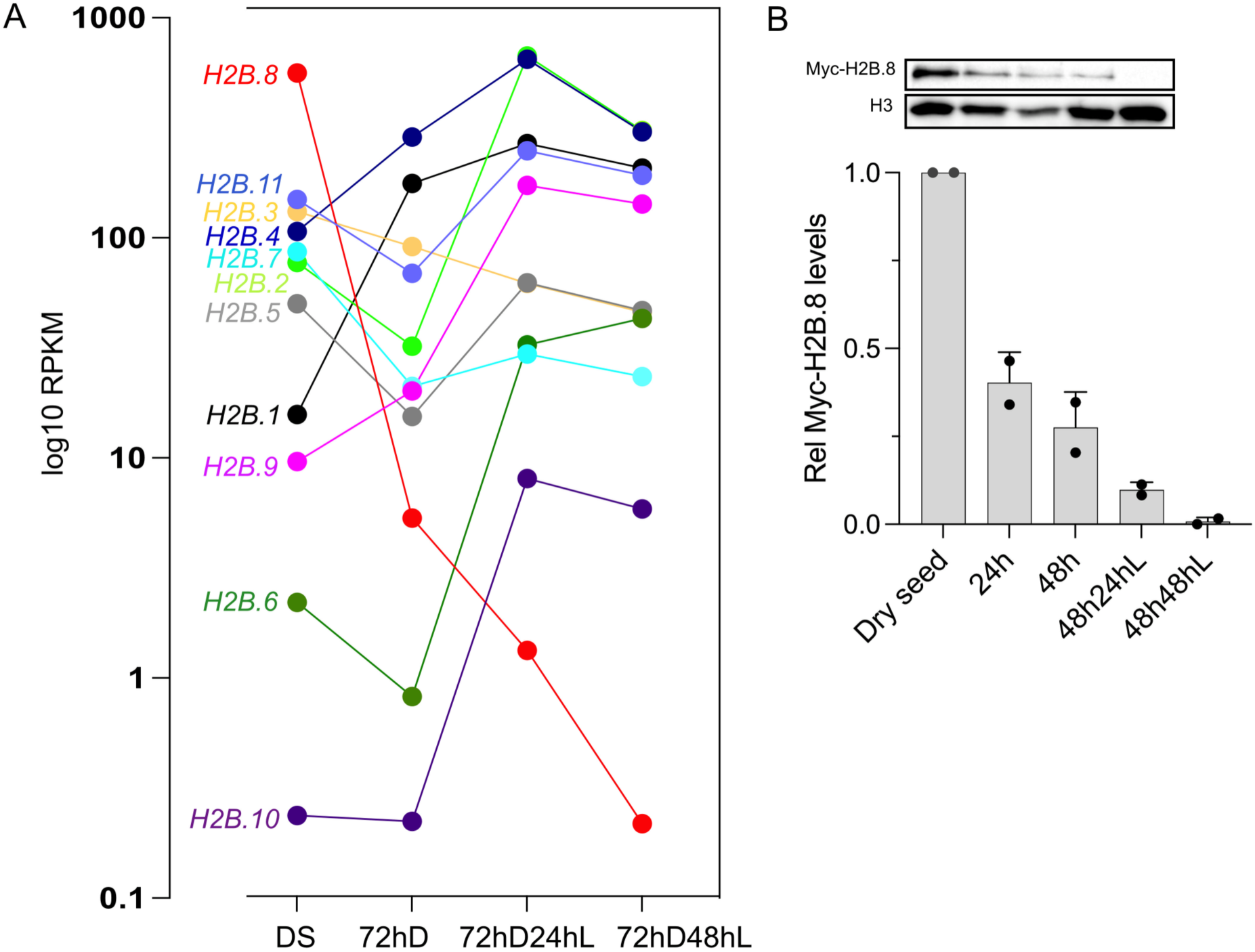
H2B.8 eviction during imbibition. **A**. Expression of the 11 *H2B* genes in DS, after 4 days of stratification in dark and cold (72hD) and 1 or 2 days after light exposure (72hD24hL, 72hD48hL, RNA-seq dataset from ^7^). **B**. Representative Western Blots showing Myc-H2B.8 and the H3 loading control (top) and quantification (bottom) of Myc-tagged H2B.8 expressed from the endogenous promoter during the seed-to-seedling transition normalized to H3. Normalized Myc-H2B.8 levels in DS were set to 1.

**Supplementary Figure 12:**
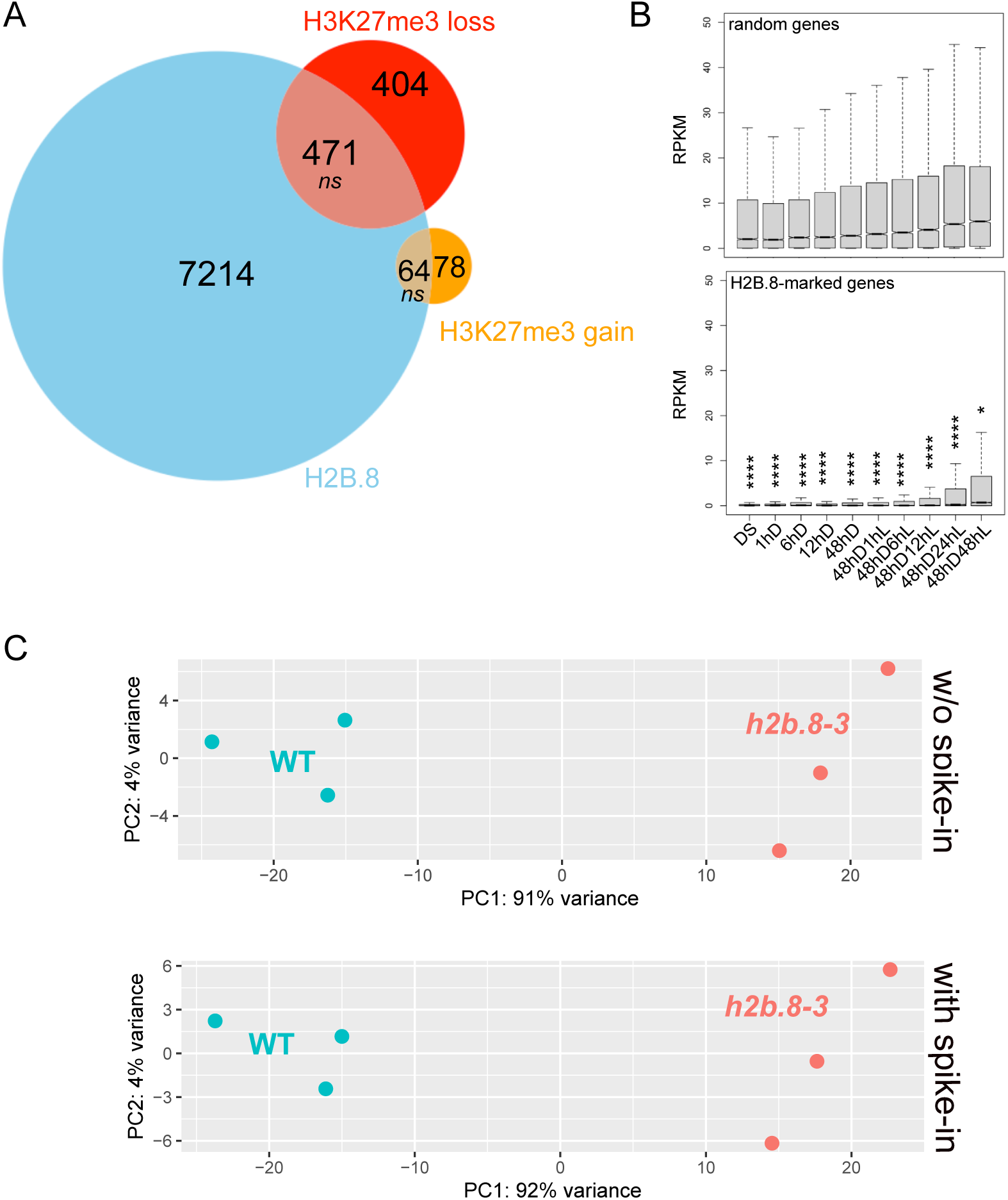
Expression of H2B.8-marked genes during the seed-to-seedling transition. **A**. Venn diagram showing overlap of H2B.8-marked genes and genes that loose or gain H3K27me3 during seed germination (48h in the light; H3K27me3 ChIP-seq datasets were obtained from ^41^); *not significant (ns)*, hypergeometric test. **B**. Differences between transcript levels of random and H2B.8-enriched genes (*\* p < 0.05, **** p< 0.0001*, non-parametric Mann-Whitney test) determined at several time points during the seed-to-seedling transition (RNA-seq data were obtained from ^8^). **C**. Principal component analysis for gene expression patterns in WT and *h2b.8-3* mutants at 24hD with and without normalization to Drosophila *spike-in* RNA.

**Supplementary Figure 13:**
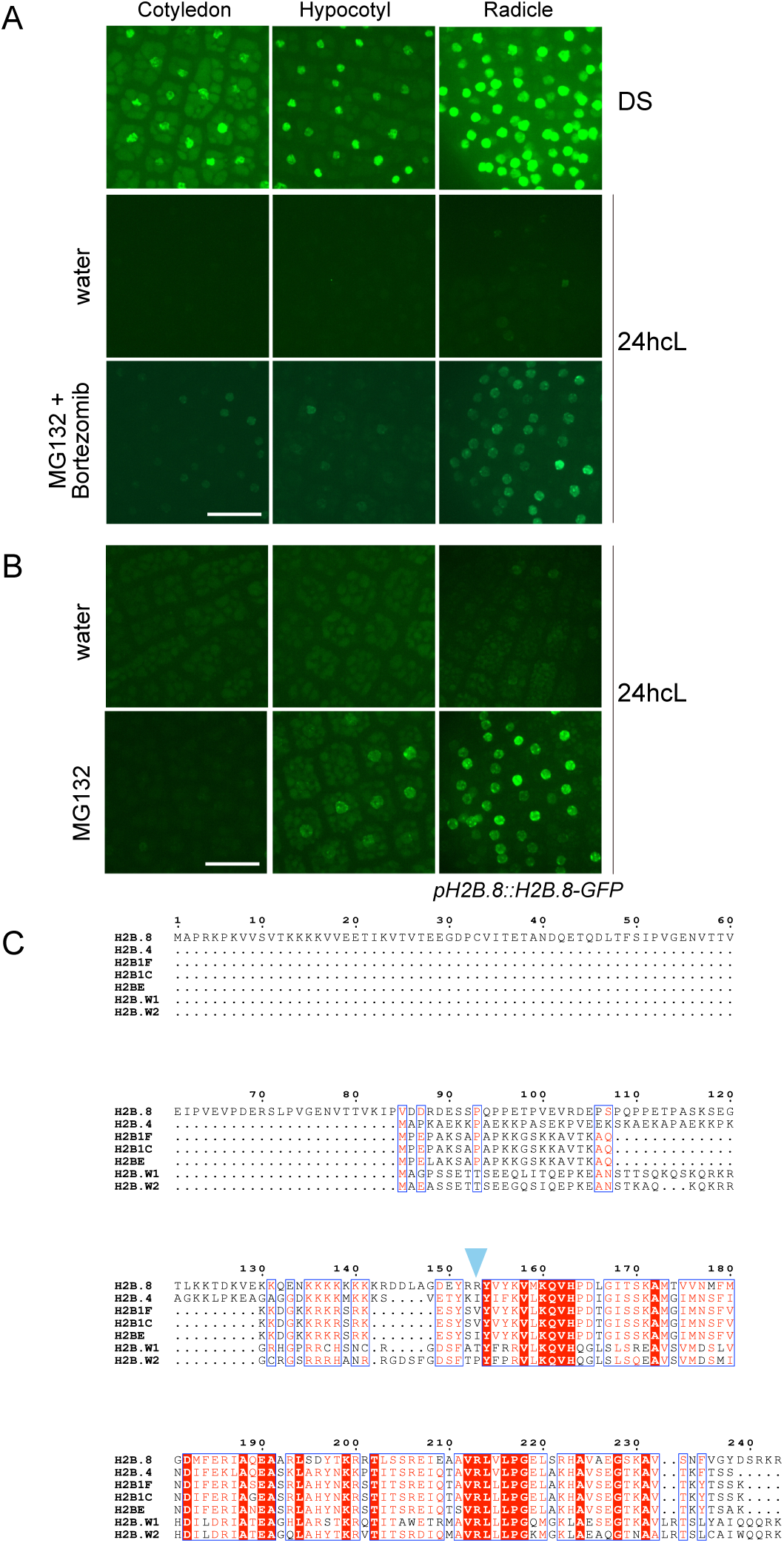
Degradation of H2B.8 during the seed-to-seedling transition A-. **B**. Representative maximum intensity projections of cotyledons, hypocotyls and radicles from embryos expressing H2B.8 as a GFP fusion protein under its endogenous promoter isolated from dry seeds or imbibed seeds under continuous light (24hcL). Images were taken with a Spinning confocal microscope, 63x objective using identical exposure settings for treated and untreated samples. Seeds were imbibed in water or in the presence of the proteasome inhibitor MG132 (100μM, **A**) or a combination of MG132 and Bortezomib (50μM each, **B**). **C**. Alignment of H2B variants from Arabidopsis (H2B.8, H2B.4), mouse (H2B1F, H2BE) and human (H2B1C, H2B.W1 and H2B.W2) generated with MUSCLE and ESPript 3.0. The blue arrowhead indicates the residue identified to alter chromatin accessibility in nucleosomes containing mouse H2BE ^54^.

## Supplementary Data

*Supplementary Data 1: Hi-C interaction pairs*

*Supplementary Data 2: Primer sequences for cloning, RT-qPCR, genotyping and ChIP-qPCR*

*Supplementary Data 3: List of RNA-, ATAC-and ChIP-seq datasets used in this study*

